# Co-targeting an AMPK–MAPK axis reprograms fibroblasts and suppresses PDAC

**DOI:** 10.64898/2026.03.16.711738

**Authors:** Ryodai Yamamura, Yusuke Satoh, Junki Fukuda, Taku Kimura, Takuya Otsuka, Sho Sekiya, Taiga Hirata, Soichiro Hata, Reo Sato, Chihiro Kamijo, Takuya Moriguchi, Shinya Kosuge, Takuya Kato, Yuya Urano, Kanako C. Hatanaka, Alexander V. Tyakht, Kazuaki Harada, Yasuyuki Kawamoto, Kazumichi Kawakubo, Masaki Kuwatani, Shintaro Takeuchi, Masataka Wada, Toshimichi Asano, Toru Nakamura, Shigeki Jin, Tomoko Mitsuhashi, Fujiko Sueishi, Kazutsune Yamagata, Atsushi Masamune, Masanobu Oshima, Takashige Abe, Nobuo Shinohara, Yoshihiro Matsuno, Yutaka Hatanaka, Shinya Tanaka, Yohei Shimono, Kotaro Matoba, Ruth E. Ley, Naoya Sakamoto, Satoshi Hirano, Tomoyoshi Soga, Shinji Fukuda, Atsushi Enomoto, Masahiro Sonoshita

## Abstract

Pancreatic ductal adenocarcinoma (PDAC) is a fatal cancer characterized by limited therapeutic options and a highly treatment-resistant tumor microenvironment. Beyond tumor-intrinsic genetic alterations, growing evidence indicates that host–microbiome interactions influence cancer progression through microbial metabolites. However, how microbiome-derived metabolites influence oncogenic signaling in PDAC remains unclear. Here, integrated profiling revealed a consistent reduction of the microbial metabolite acetic acid in fecal samples from treatment-naïve patients with PDAC and in a genetically defined *Drosophila* model recapitulating key PDAC driver alterations. Acetic acid activates AMP-activated protein kinase, and pharmacological activation of this pathway together with inhibition of mitogen-activated protein kinase signaling suppressed tumor growth in fly and mouse models. Combined pathway targeting restored AMPK activity and suppressed cancer-associated fibroblast activation. These findings identify a microbiome-associated metabolic vulnerability in PDAC and suggest that coordinated targeting of metabolic and oncogenic signaling may restrain tumor progression and improve therapeutic strategies.

## INTRODUCTION

Pancreatic ductal adenocarcinoma (PDAC) is projected to become a leading cause of cancer-related mortality globally. Consistent with this, the 5-year overall survival rate of PDAC remains poor at approximately 13% across all disease stages^1^. Current standard therapies, including surgical resection and combination chemotherapies such as FOLFIRINOX and gemcitabine with nab-paclitaxel, provide modest survival benefits but are frequently associated with significant toxicity. Moreover, emerging immunotherapeutic strategies, including immune checkpoint blockade and chimeric antigen receptor T cell therapies, have shown limited efficacy in PDAC, largely owing to the profoundly immunosuppressive tumor microenvironment. Although targeted and combination approaches are under active investigation, durable clinical responses remain limited and therapeutic resistance frequently emerges. Collectively, these limitations underscore an urgent need to identify fundamentally novel therapeutic vulnerabilities and treatment paradigms for PDAC.

Recent studies have identified the gut microbiome as an important modulator of tumor progression and treatment response across multiple types of cancer, including PDAC, highlighting its potential as a source of novel therapeutic targets^2,3^. Although the underlying mechanisms remain incompletely understood, accumulating evidence suggests that microbiome-derived metabolites play central roles in shaping tumor biology. For example, bile acids modified by gut bacteria can induce cyclooxygenase-2 (COX-2) in host tissues, thereby influencing immune regulation^4^. In addition, short-chain fatty acids (SCFAs), a major class of microbiome-derived metabolites, have been reported to attenuate tumor-associated immunosuppression and enhance host anti-tumor immune responses, being correlated with improved prognosis and enhanced chemotherapy responsiveness in PDAC^5,6^. Together, these findings are consistent with an emerging view that microbial functional output, rather than the mere presence or absence of individual taxa, contributes to the regulation of tumor behavior, therapeutic responsiveness, and the immune landscape in PDAC.

Building on these insights, systematic interrogation of microbiome–tumor–host interactions in PDAC, with a particular focus on gut microbiome-derived metabolites as functional mediators, represents a key frontier in the field. However, previous studies have often been constrained by small cohort sizes, limited taxonomic resolution or suboptimal matching of healthy control populations, leaving microbiome–metabolite associations in PDAC incompletely defined. To address these limitations, we established an integrated multi-omics framework that combines well-characterized patient-derived samples with complementary whole-animal models, enabling mechanistic interrogation of microbiome-associated metabolic vulnerabilities in PDAC.

## RESULTS

### Gut microbiome dysbiosis and reduced microbial acetic acid (AA) production in patients with PDAC

To determine whether PDAC is associated with alterations in gut microbiome composition and microbial metabolic output, we performed integrated microbiome and metabolic profiling of fecal samples from 33 treatment-naïve patients with PDAC and 42 healthy controls (HCs), as outlined in Figure 1A and Table S1.

**Figure 1.**
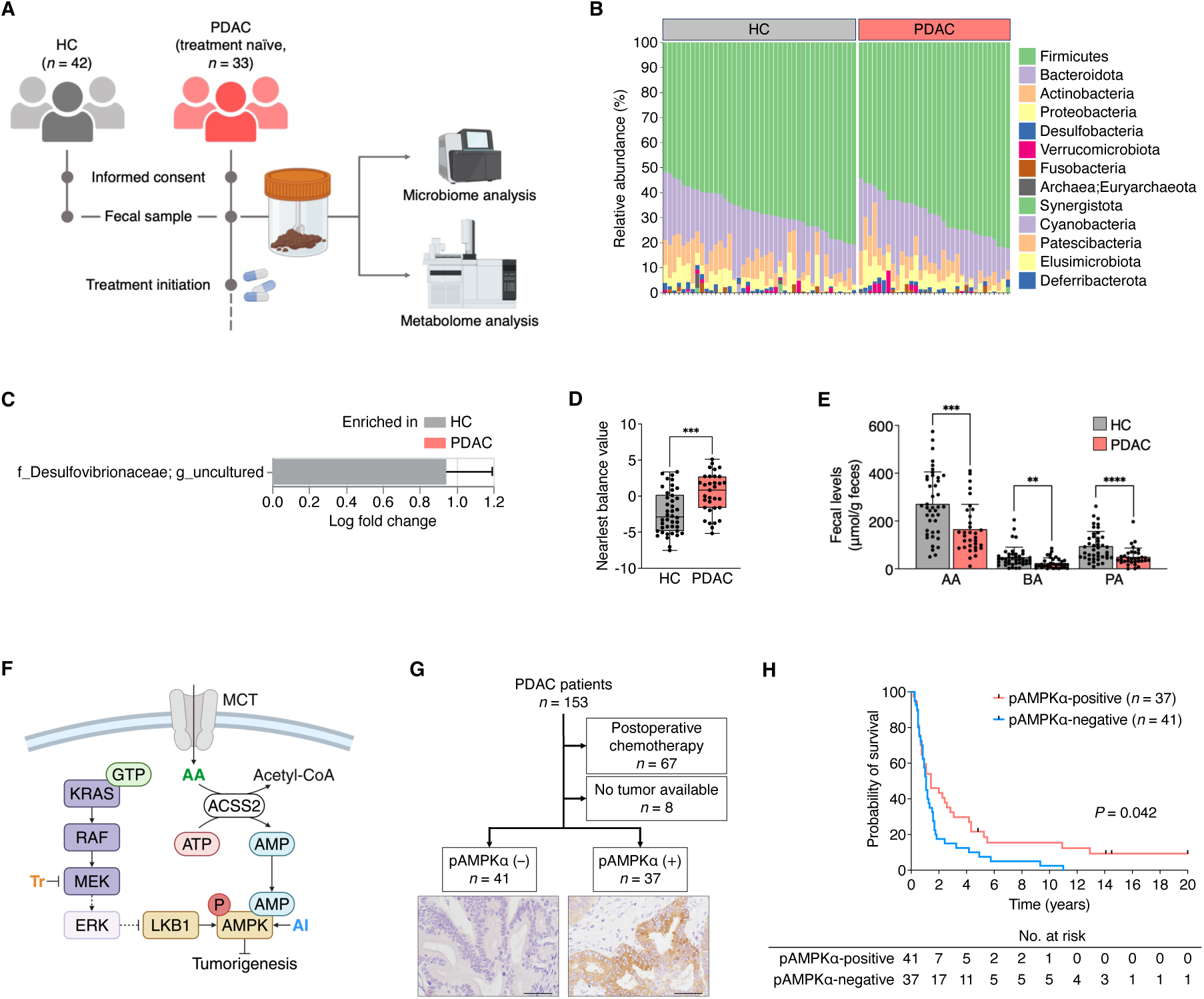
Gut microbiome dysbiosis, reduced microbial AA production and impaired AMPK signaling in patients with PDAC. **A,** Schematic overview of the clinical study. Fecal samples were collected from 42 HCs and 33 treatment-naïve patients with PDAC and subjected to 16S rRNA sequencing for microbiome profiling and GC/MS for SCFA quantification. **B,** Relative abundance of bacterial phyla in the HC and PDAC groups. **C,** Reduced abundance of Desulfovibrionaceae in patients with PDAC, as assessed by ANCOM-BC. **D,** NB values are significantly elevated in patients with PDAC relative to HCs, indicating a PDAC-associated shift in gut microbiome composition. Boxes indicate the interquartile range (IQR), and the line represents the median. **E,** Fecal levels of major SCFAs, including AA, BA and PA, are significantly lower in patients with PDAC than in HCs, as quantified by GC/MS. **F,** Schematic model illustrating intracellular AA metabolism and its downstream regulation of the AMPK pathway. AA is accumulated via monocarboxylate transporters (MCTs) and converted to acetyl-CoA by cytoplasmic ACSS2, a reaction that consumes ATP and generates AMP. The resulting increase in the AMP/ATP ratio promotes binding of AMP to the γ subunit of AMPK, inducing an allosteric conformational change that facilitates phosphorylation of the α subunit at Thr172 by LKB1, a principal upstream AMPK kinase, thereby activating AMPK. In parallel, the AMPK activator AI is accumulated by cells and phosphorylated to ZMP, an AMP mimetic that directly activates AMPK by promoting γ subunit binding independently of changes in the cellular AMP/ATP ratio. Activated AMPK can in turn suppress RAF activity, establishing a feedback loop between AMPK and the RAF–MAPK signaling axis. Conversely, the activated MAPK pathway suppresses AMPK signaling through ERK-mediated inhibitory phosphorylation of LKB1 at Ser428. MEK inhibition by Tr might relieve this inhibitory modification and restore LKB1–AMPK coupling, enabling cooperative AMPK activation with AA-driven metabolic signaling. **G,** Patient-selection scheme for survival analysis, with representative immunohistochemical images of pAMPKα staining in human PDAC tumors. Of the 153 PDAC cases initially included in the TMAs, 67 patients who received postoperative chemotherapy and 8 samples lacking evaluable tumor tissue were excluded, yielding 78 evaluable cases for final analysis. Patients were stratified into pAMPKα-negative (*n* = 41) and pAMPKα-positive (*n* = 37) groups. Scale bars, 50 μm. **H,** Kaplan–Meier analysis demonstrating significantly longer overall survival in patients with pAMPKα-positive PDAC than in pAMPKα-negative patients. Numbers at risk are presented below the plot. Each dot represents an individual human subject. Statistical significance was assessed using a two-sided Student’s *t*-test (**D**), one-way ANOVA with Tukey’s multiple comparisons test (**E**), and the log-rank test for survival analysis (**H**). \*\**P* < 0.01, \*\*\**P* < 0.001, \*\*\*\**P* < 0.0001. Data in **C** are presented as the mean ± SEM; data in **D** are presented as the median and IQR, with whiskers indicating 1.5 × IQR; data in **e** are presented as the mean ± SD.

At the phylum level, no substantial differences in microbiome diversity were observed between the groups, and overall community diversity did not significantly differ (beta-diversity analysis; Figure 1B, Figure S1A). Conversely, analysis of compositions of microbiomes with bias correction (ANCOM-BC) revealed significant depletion of uncultured taxa (assigned to the Desulfovibrionaceae family) in PDAC samples relative to HC samples (*q* = 0.018; Figure 1C, Figure S1B). Members of this family have been reported to produce AA by oxidizing organic substrates^7^. Importantly, this association remained statistically significant after adjustment for potential confounding factors using multi-variable association with linear models 2 (MaAsLin2), including models adjusted for age and sex, as well as those further adjusted for body mass index (BMI) and type 2 diabetes mellitus (T2DM) status (Figure S1C, D).

To further define PDAC-associated microbial compositional features beyond individual taxa, we applied a compositionally aware Nearest Balance (NB) analysis^8^. This approach summarises microbiome variation between the groups using a ratio between two sets of taxa, thereby mitigating compositional bias inherent to relative abundance data and capturing global shifts in community structure. Based on the marginal association between general microbiome composition at the genus level and clinical status, as determined by permutational multivariate analysis of variance (permutational analysis of variance (PERMANOVA) over weighted UniFrac distance, *P* = 0.080), the obtained PDAC-associated genus-level NB in its denominator (i.e. among the taxa enriched in HC) included *Phascolarctobacterium*, *Collinsella*, *Anaerostipes*, *Fusicatenibacter*, *Sutterella*, and *Faecalibacterium* (Figure S1E). Conversely, the numerator (i.e. taxa enriched in PDAC) included *Alistipes*, *Streptococcus* and *Odoribacter* (Figure S1E). In line with NB definition, the distribution of per-sample NB values was significantly higher for PDAC patients than for HC subjects (Figure 1D), indicating a global shift in microbial composition favoring PDAC-associated taxa.

Given these PDAC-associated alterations in microbial community structure, we next examined whether corresponding changes existed in microbiome-derived metabolites by performing targeted metabolomic profiling of fecal samples, focusing on SCFAs, which are important metabolic products of the gut microbiome and key regulators of host immunity and tumor progression^9,10^. Specifically, we quantified nine microbiota-related organic acids, including the canonical SCFAs AA, propionic acid (PA), butyric acid (BA), isobutyric acid (IBA), isovaleric acid (IVA) and valeric acid (VA) as well as lactic acid (LA), formic acid (FA) and succinic acid (SA), using gas chromatography–mass spectrometry (GC/MS). Among these, fecal concentrations of AA, BA and PA were significantly lower in patients with PDAC than in HCs (Figure 1E, Table S2). These differences remained statistically significant after adjusting for age, sex, BMI and T2DM status (Table S2). Given that intestinal SCFAs, specifically AA, are almost exclusively derived from microbial fermentation of dietary fiber^11^, the observed reductions in fecal AA levels in patients with PDAC likely reflect a functional deficit in the gut microbiome. Furthermore, the persistence of these metabolic differences after adjusting for clinical covariates suggests that this reduced AA availability is an inherent feature of the PDAC-associated state.

Collectively, these results indicate that PDAC is associated with a compositional shift in the gut microbiome, characterized by relative depletion of members of the *Desulfovibrionaceae* family and other commensal genera, such as *Collinsella* and *Phascolarctobacterium*, which have been reported to contribute to AA production^7,12,13^, together with enrichment of taxa including *Alistipes*, *Streptococcus* and *Odoribacter*. This microbial dysbiosis was accompanied by a coordinated reduction in fecal levels of key SCFAs. Notably, among the organic acids measured, AA exhibited the most pronounced reduction in patients with PDAC and remained robustly significant even after stringent adjustment for major clinical covariates (Table S2). Although these findings do not establish causality, they are consistent with a model in which microbiome alterations favoring PDAC-associated taxa are linked to reduced AA availability, a prominent metabolic feature of the PDAC-associated condition.

### AMP-activated protein kinase (AMPK) activation correlates with longer survival in patients with PDAC

To explore potential signaling pathways that may link microbiome-derived AA to tumor behavior in PDAC, we focused on AMPK, a central metabolic sensor that integrates cellular energy status with growth control. *In vivo*, AA derived from the diet or produced by gut microbes is absorbed by colonocytes and rapidly converted to acetyl-CoA by cytoplasmic acetyl-CoA synthetase 2 (ACSS2)^14^. This ATP-consuming reaction increases the intracellular AMP/ATP ratio, thereby activating AMPK through allosteric AMP binding to the γ subunit and phosphorylation of the catalytic α subunit at Thr172 by upstream kinases such as liver kinase B1 (LKB1)^15^ (Figure 1F).

Previous studies also demonstrated extensive cross-talk between the AMPK and mitogen-activated protein kinase (MAPK) pathways. Activated AMPK suppresses tumorigenesis through multiple mechanisms, including inhibition of BRAF^16^ signaling downstream of RAS^16^, activation of p53^17^, suppression of mammalian target of rapamycin signaling via tuberous sclerosis complex 2^18^ and repression of aerobic glycolysis^19^ (Figure 1F). Conversely, oncogenic BRAF^V600E^ activates extracellular signal-regulated kinase (ERK) and p90RSK, leading to inhibitory phosphorylation of LKB1 and functional uncoupling of the LKB1–AMPK axis^20,21^. Notably, pharmacological inhibition of BRAF^V600E^ has been shown to restore AMPK activation^22^ (Figure 1F).

Based on our observation that fecal AA concentrations are reduced in PDAC patients, we hypothesised that the decrease in luminal AA availability, which potentially reflects the identified microbial shifts as well as altered production or uptake rates, may attenuate AMPK activation and contribute to tumor progression. To assess the clinical relevance of AMPK activation in PDAC, we examined tumor tissues using tissue microarrays (TMAs) from a patient cohort that was distinct from the fecal microbiome analysis cohort. Because AMPK activation critically depends on phosphorylation of its α subunit within the kinase domain, we focused on phosphorylated AMPKα (pAMPKα) as a marker of pathway activation. Immunohistochemical analysis of 78 evaluable PDAC specimens revealed detectable pAMPKα staining in cancer cells in 37 specimens (47.4%; Figure 1G, Table S3). Patients with pAMPKα-positive tumors exhibited significantly longer overall survival than patients with negative staining (Figure 1H), and this association remained significant after adjustment for clinical covariates including age and sex (Table S4). To determine whether this prognostic association reflected AMPK activation rather than overall protein abundance, we additionally assessed total AMPKα (tAMPKα) expression in the same cohort. Although 39 of 78 patients (50.0%) were classified as tAMPKα-high based on the median H-score, tAMPKα expression showed no significant association with overall survival (Figure S1F, G, Table S5, 6).

Collectively, these data indicate that phosphorylation-dependent activation of AMPK, rather than total AMPKα abundance, is associated with favorable clinical outcomes in PDAC. These findings identify AMPK activation as a clinically relevant feature of PDAC and provide a rationale for therapeutic strategies aimed at restoring or enhancing AMPK signaling in this malignancy.

### PDAC-associated oncogenic alterations drive gut microbiome dysbiosis and reduced AA abundance in Drosophila

To investigate the biological significance and potential causal contribution of the microbiome and metabolite alterations observed in patients with PDAC, we employed a genetically tractable whole-organism model. *Drosophila melanogaster* provides a powerful experimental system for dissecting complex interactions among oncogenic signaling, host metabolism and the tissue microenvironment, with growing recognition of its translational relevance to human cancer biology^23–27^. Fly models have been particularly informative in uncovering how oncogenic pathways interface with organismal physiology to shape malignant phenotypes, including cell competition, metabolic re-programming and diet-dependent tumor growth^28,29^.

In this study, we employed a previously established *Drosophila* model (*Serrate* {*Ser*}-*gal4*;*UAS*-*Ras*^G12D^,*UAS*-*p53*^shRNA^,*UAS*-*dCycE*,*UAS*-*Med*^shRNA^, hereafter referred to as *4-hit*) that recapitulates four genetic alterations associated with the worst prognosis in patients with PDAC–*KRAS*^G12D^, *TP53* loss-of-function (LOF), *CDKN2A* LOF and *SMAD4* LOF^30–33^. Although *Drosophila* does not develop *bona fide* carcinomas, this model exhibits profound epithelial transformation, disrupted tissue architecture and organismal lethality, thereby capturing key oncogenic features relevant to aggressive PDAC biology. Leveraging this platform, we have previously identified multiple therapeutic vulnerabilities that synergise with MEK inhibition to suppress oncogenic phenotypes, including aurora kinase B, glycogen synthase kinase 3 and the nicotinamide adenine dinucleotide–glutathione peroxidase 4 axis^30–32^.

We first investigated whether PDAC-associated oncogenic signaling is sufficient to influence gut microbiome composition in this model. Comparison of the gut microbiome between *4-hit* flies and genetic controls (*Ser*-*gal4* × *white*^−^) revealed no major differences at the phylum level (Figure 2A, B). Consistently, overall community structure did not differ significantly between the two groups based on beta-diversity analysis using weighted UniFrac distances (as determined by PERMANOVA, *P* = 0.117; Figure S2A). Despite the absence of global community shifts, taxon-specific differences were detected by ANCOM-BC, which identified a marked depletion of *Acetobacter* in *4-hit* flies (*q* = 0.00002). *Acetobacter* is a well-characterized commensal of *Drosophila* and a major microbial source of AA^34^. This depletion was accompanied by enrichment of *Methylobacterium/Methylorubrum* (*q* = 0.015) and GCA-900066575 (*q* = 0.024) (Figure 2C, Figure S2B). At the species level, *Acetobacter* reads were most closely aligned with *Acetobacter persici* (AP).

**Figure 2.**
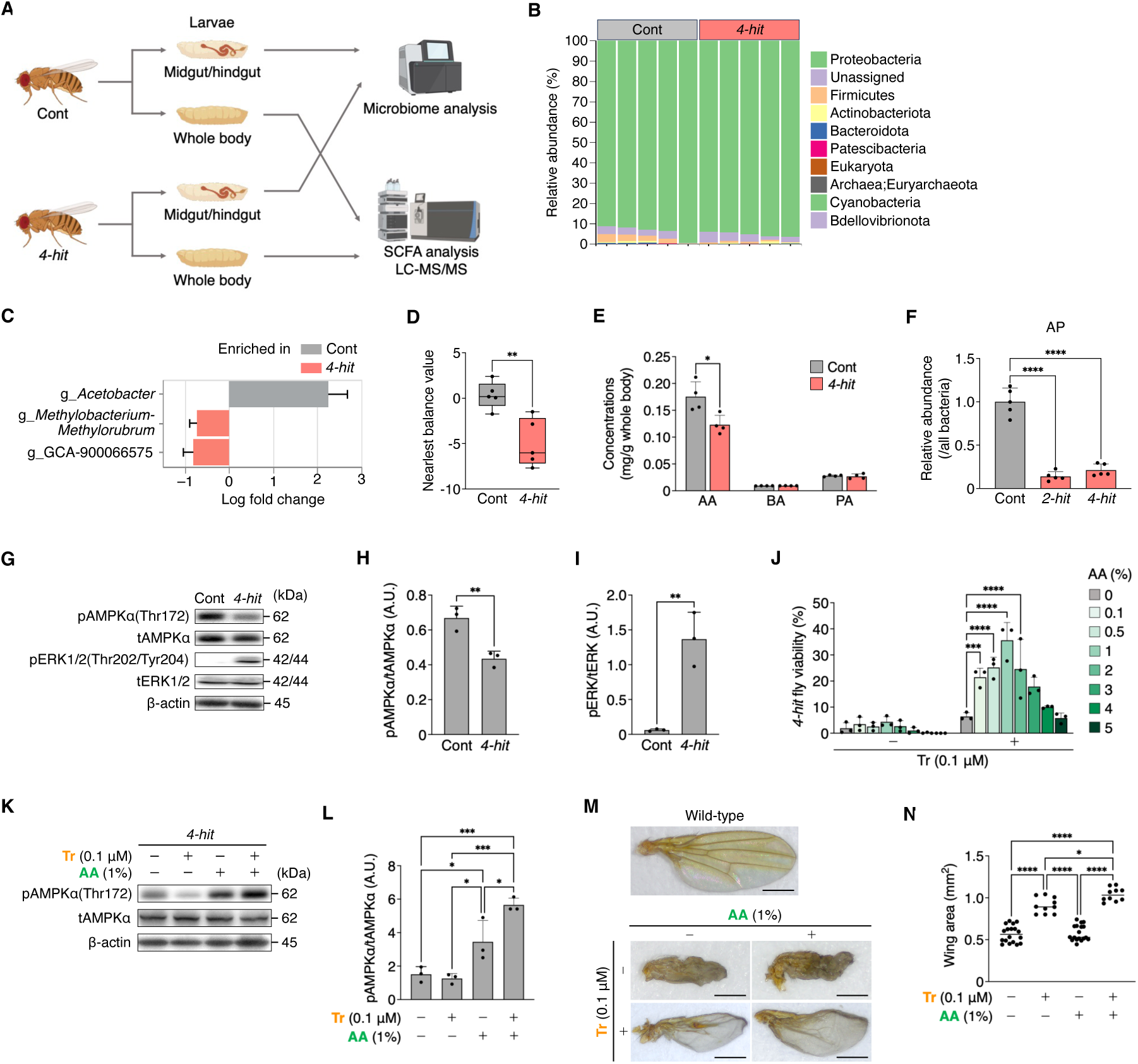
Gut microbiome dysbiosis and reduced AA production enable synergistic MAPK–AMPK targeting in a *Drosophila* model of PDAC-associated key genetic abnormalities. **A,** Experimental workflow for microbiome and metabolome analyses in *Drosophila*. Control (Cont; *Serrate* {*Ser*}-*gal4* × *white*^−^) and *4-hit* larvae, recapitulating combined *KRAS–TP53–CDKN2A–SMAD4* alterations, were subjected to 16S rRNA sequencing of midgut and hindgut samples and LC-MS/MS-based quantification of SCFAs in whole larvae. **B,** Relative abundance of bacterial phyla in control (*n* = 5) and *4-hit* (*n* = 5) larvae. **C,** ANCOM-BC of midgut and hindgut microbiota reveals a distinct gut microbial composition in *4-hit* larvae, characterized by marked depletion of *Acetobacter* compared with controls. **D,** NB analysis reveals a significant shift in microbiome composition in *4-hit* larvae away from taxa enriched in controls, reflected by reduced NB values. Boxes indicate the interquartile range (IQR), and the center line represents the median. **E,** Whole-body concentrations of AA, BA and PA in control and *4-hit* larvae quantified by LC-MS/MS. **F,** Colonization by the AA-producing bacterium AP is markedly impaired in flies harboring oncogenic genetic alterations. Relative AP abundance was quantified by qPCR in control, *2-hit* larvae (recapitulating combined *KRAS-TP53* alterations) and *4-hit* larvae. G–I, *4-hit* larvae exhibit reduced phosphorylation of AMPKα and elevated phosphorylation of ERK. **G,** Immunoblot analysis of phosphorylated and total AMPKα and ERK in control and *4-hit* larvae. β-actin served as a loading control. **H, I,** Quantification of the ratios of phosphorylated to total AMPKα (pAMPKα/tAMPKα) (**H**) and ERK (pERK/tERK) (**I**). **J,** AA synergises with Tr (0.1 μM) to enhance the viability of *4-hit* flies. Compound concentrations indicate final concentrations in fly food. **K–L,** Combined Tr+AA treatment enhances AMPKα phosphorylation. **K,** Immunoblotting of AMPKα phosphorylation in *4-hit* larvae treated with Tr (0.1 μM) and/or AA (1%). β-actin served as a loading control. **L,** Quantification of the pAMPKα/tAMPKα ratio. **M–N,** Tr+AA suppresses wing deformities and increases wing size in *4-hit* flies. **M,** Representative images of adult wings from wild-type and *4-hit* flies treated with AA and/or Tr. Scale bars, 500 μm. **N,** Quantification of the adult wing area. Each dot represents an individual fly wing. Each dot represents an individual sample consisting of pooled midgut and hindgut tissues from 10 fly larvae (**D**), an individual sample consisting of 50 mg of fly larval tissue (**E**), an independent experiment (**F, H–J, L**) or an individual fly wing (**N**). Statistical significance was assessed by a two-sided Student’s *t*-test (**D–E, H–I**) and one-way ANOVA followed by Tukey’s multiple comparisons test (**F, L, N**) or Dunnett’s test (**J**). \**P* < 0.05, \*\**P* < 0.01, \*\*\**P* < 0.001, \*\*\*\**P* < 0.0001. Data in **D** are presented as the median with the IQR, with whiskers indicating 1.5 × IQR; all other data are presented as the mean ± SD. A.U., arbitrary units.

To further characterize subtle taxonomic shifts, we performed a compositional-aware NB analysis, using the same approach as in the human cohorts. Although this analysis corroborated the absence of large-scale community restructuring, genus-level NB analysis revealed that denominator taxa, tentatively enriched in control flies, comprised *Acetobacter* and *Megamonas*, whereas numerator taxa, tentatively enriched in *4-hit* flies, included *Blautia*, *Acinetobacter*, *Bacteroides*, and *Streptococcus* (Figure S2C). Consistent with these patterns, NB values were significantly lower in *4-hit* flies than in controls (Figure 2D), suggesting that *Acetobacter* is selectively disadvantaged in the *4-hit* genetic context.

We next examined whether the reduced abundance of AA-producing bacteria was accompanied by altered metabolite levels. Quantitative PCR (qPCR) confirmed a reduction in AP abundance in *4-hit* flies (Figure S2D). Consistently, liquid chromatography–tandem mass spectrometry (LC-MS/MS) analysis revealed a significant decrease in whole-body AA levels in *4-hit* larvae, whereas levels of BA and PA were not significantly altered (Figure 2E).

Having observed reductions in major AA-producing bacterial taxa together with decreased fecal and systemic AA levels in both patients with PDAC and *4-hit* flies, we next examined whether oncogenic alterations influence bacterial colonization capacity. We hypothesised that oncogenic signaling may generate a host microenvironment that is less permissive to stable colonization by specific commensal taxa. To test this, we orally administered AP to control flies, *2-hit* flies mimicking combined *RAS*–*p53* alterations and *4-hit* flies and subsequently quantified colonization efficiency. AP colonization was significantly reduced in both *2-hit* and *4-hit* flies compared with controls, with no appreciable difference between the two mutant genotypes (Figure 2F).

Together, these findings indicate that PDAC-relevant oncogenic genetic alterations are sufficient to impair colonization of the fly gut by AA-producing bacteria in a whole-organism model. This host genotype-dependent constraint on microbial colonization provides evidence for a mechanistic connection between oncogenic signaling, gut microbiome dysbiosis, and reduced AA availability in PDAC-associated disease contexts.

### Trametinib (Tr) and AA exhibit synergistic effects

To determine whether the reduction in AA levels observed in PDAC-associated oncogenic states is linked to impaired AMPK activation *in vivo*, we first compared AMPK signaling between control and *4-hit* flies. Immunoblot analysis revealed a significant reduction in AMPKα phosphorylation in *4-hit* flies relative to controls, accompanied by a pronounced increase in ERK phosphorylation (Figure 2G–I). This reciprocal pattern is consistent with oncogenic MAPK signaling imposing inhibitory constraints on the LKB1–AMPK axis (Figure 1F), leading us to hypothesise that concurrent inhibition of MAPK signaling and activation of AMPK might restore AMPK function and ameliorate the lethal phenotype of *4-hit* flies.

To test this hypothesis, we co-administered AA with Tr, a clinically approved MEK inhibitor used to treat *BRAF*-mutant malignancies, including melanoma and non-small-cell lung cancer, and previously shown to improve the viability of *4-hit* flies^30–32^. Co-treatment with AA (0.1%–2% in fly food) significantly enhanced the viability-rescuing effect of Tr compared with Tr monotherapy (Figure 2J). Quantitative synergy analysis using the zero interaction potency (ZIP) model identified the combination of 0.1 μM Tr and 1% AA as the most synergistic across all tested conditions (Figure S3A). Consistent with these phenotypic effects, immunoblotting confirmed that combined Tr and AA treatment robustly increased AMPKα phosphorylation in *4-hit* larvae (Figure 2K, L). Although AA alone induced a measurable increase in AMPKα phosphorylation in control larvae and a trend toward increased phosphorylation in *4-hit* larvae (Figure S3B–E), this activation was insufficient to confer a survival benefit. These findings indicate that AMPK activation is functionally constrained in the presence of persistent oncogenic MAPK signaling, and that effective phenotypic rescue requires concomitant relief of MAPK-mediated suppression.

We next sought independent *in vivo* validation of the functional interplay between MAPK and AMPK signaling using adult wing morphology, a well-established *Drosophila* readout that is highly sensitive to RAS pathway activity^30–32^. Although *4-hit* flies raised at 16°C remain viable because of reduced GAL4-driven transgene expression and attenuated cellular transformation, untreated flies nonetheless exhibited pronounced wing vein thickening and structural deformities (Figure 2M), consistent with prior observations^30–32^. Combined administration of Tr and AA (Tr+AA) markedly suppressed these abnormalities, producing a substantially stronger corrective effect than either treatment alone (Figure 2M, N). This tissue-level phenotypic normalization further supports the notion that simultaneous modulation of MAPK and AMPK signaling counteracts RAS-driven pathological outputs *in vivo*.

Collectively, these findings demonstrate that Tr and AA act synergistically in a whole-organism model to restore AMPK activation and suppress oncogenic phenotypes. Together, they support a therapeutic framework in which coordinated targeting of the MAPK and AMPK pathways may provide enhanced control over PDAC-associated oncogenic programs.

### Genetic modulation of AMPK signaling influences 4-hit Drosophila phenotypes

To obtain genetic evidence that the AMPK pathway functions as a determinant of oncogenic fitness in PDAC-associated contexts, we next performed targeted genetic modulation experiments in *4-hit* flies. We focused on core components of the AMPK pathway, including *AMPKα* (the *Drosophila* orthologue of human *PRKAA1*), *SNF4Aγ* (*PRKAG1*), *Lkb1* (*LKB1*) and *AcCoAS* (*ACSS2*), all of which play central roles in cellular energy sensing and AA metabolism. Using previously established genetic approaches^30–32^, we introduced heterozygous LOF alleles of each gene into the *4-hit* background and assessed organismal viability. Heterozygosity of any of these AMPK pathways components significantly reduced the viability of *4-hit* flies (Figure 3A), consistent with our pharmacological findings. These results indicate that intact systemic AMPK pathway activity is required to sustain organismal fitness under oncogenic stress.

**Figure 3.**
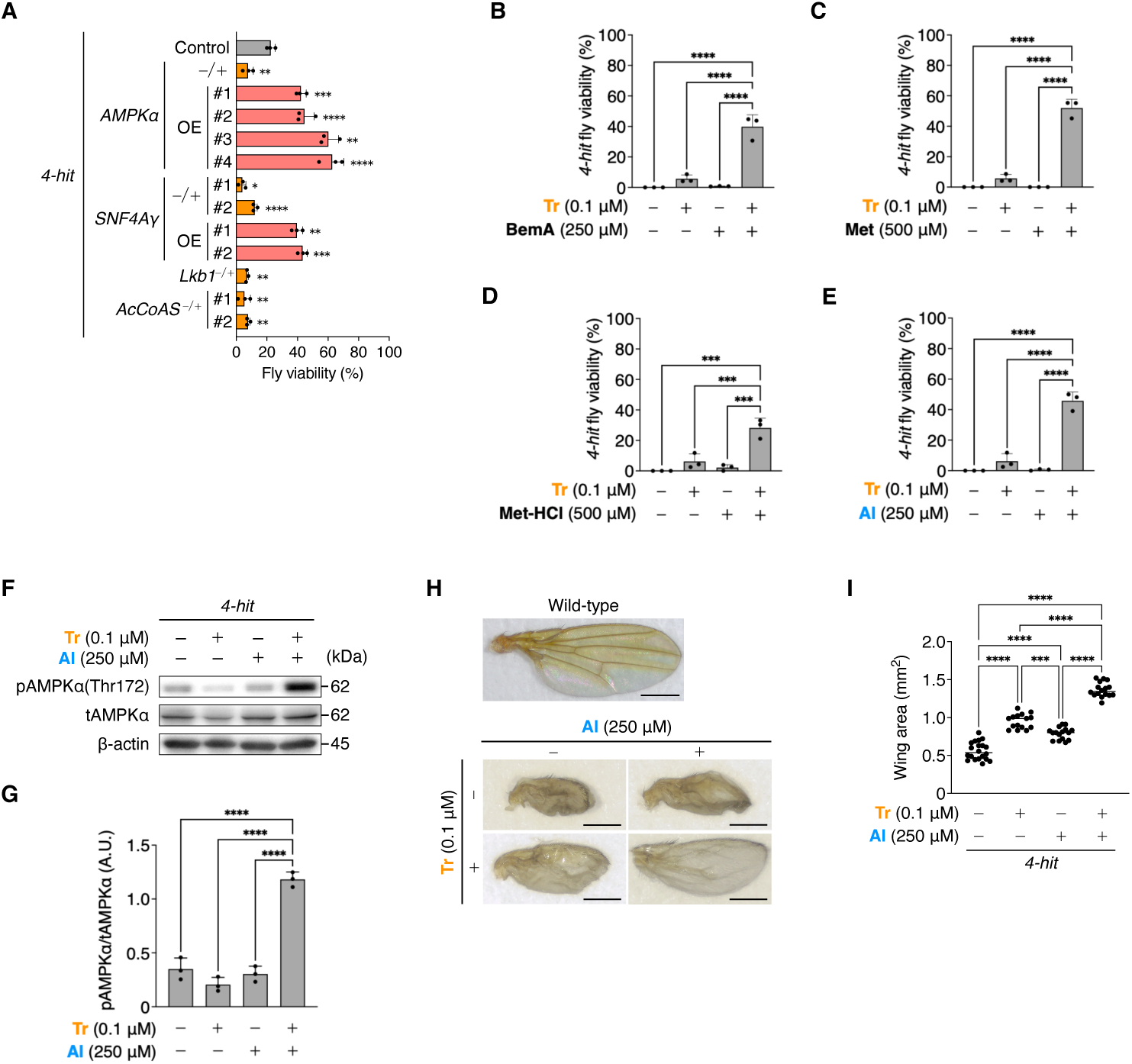
AMPK activators synergise with Tr to suppress transgene-induced abnormalities in *4-hit* flies. **A,** Viability of *4-hit* flies following genetic manipulation of AMPK pathway components. Partial LOF using heterozygous mutations (−/+) in *AMPKα* (the *Drosophila* orthologue of human *PRKAA1*), *SNF4Aγ* (*PRKAG1*), *Lkb1* (*LKB1*) and *AcCoAS* (*ACSS2*). In parallel, transformed-cell-specific over-expression of *AMPKα* and *SNF4Aγ* was evaluated. Viability is expressed relative to control flies. **B–E,** Four AMPK activators exhibit significant synergistic effects with Tr (0.1 μM) in rescuing lethality of *4-hit* flies. The compounds tested were BemA (250 μM) (**B**), Met (500 μM) (**C**), Met-HCl (500 μM) (**D**) and AI (250 μM) (**E**). **F–G,** Combined treatment with Tr and AI significantly enhances AMPKα phosphorylation. **F,** Immunoblotting of AMPKα phosphorylation in *4-hit* larvae treated with Tr (0.1 μM) and/or AI (250 μM). β-actin served as a loading control. **G,** Quantification of the pAMPKα/tAMPKα ratio. A.U., arbitrary units. **H–I,** Tr and AI synergistically suppress wing deformities and increase wing size in *4-hit* flies. **H,** Representative adult wings of wild-type and *4-hit* flies treated with Tr (0.1 μM) and/or AI (250 μM). Scale bars, 500 μm. **I,** Quantification of the adult wing area. Each dot represents an independent experiment (**A–E, G**) or an individual fly wing (**I**). Statistical significance was assessed by one-way ANOVA followed by Dunnett’s test (**A**) or Tukey’s multiple comparisons test (**B–E, G, I**). \*\**P* < 0.01, \*\*\**P* < 0.001, \*\*\*\**P* < 0.0001. Data are presented as the mean ± SD.

To distinguish systemic effects from cell-autonomous contributions within transformed cells, we next manipulated AMPK pathway genes specifically in transformed cells using GAL4-dependent *UAS*-driven over-expression (*UAS*-*gene*^OE^) or RNAi-mediated knockdown. Notably, transformed cell-specific over-expression of *AMPKα* or *SNF4Aγ* significantly rescued the lethality of *4-hit* flies, whereas targeted knockdown of these components further exacerbated lethality (Figure 3A).

Together, these genetic data provide independent and complementary evidence that AMPK signaling exerts a tumor-suppressive role in the *4-hit* model, acting at least in part through cell-autonomous mechanisms. These findings further substantiate the rationale for therapeutically targeting AMPK signaling in PDAC-associated oncogenic programs.

### Tr and AMPK activators exhibit synergistic effects

Although AA served as a useful mechanistic probe, its direct therapeutic application in humans is limited. Orally administered AA is rapidly absorbed and extensively metabolised in the intestinal epithelium and liver, resulting in poor pharmacokinetic (PK) controllability. Moreover, high local concentrations of AA in the intestine can induce gastrointestinal toxicity and robust inflammatory responses in mammals, including colitis-like pathology, mucosal ulceration and lethality^35,36^ (see Figure 4B, C, Figure S5A, G). Together, these PK and safety considerations preclude the direct clinical application of AA as a therapeutic agent.

**Figure 4.**
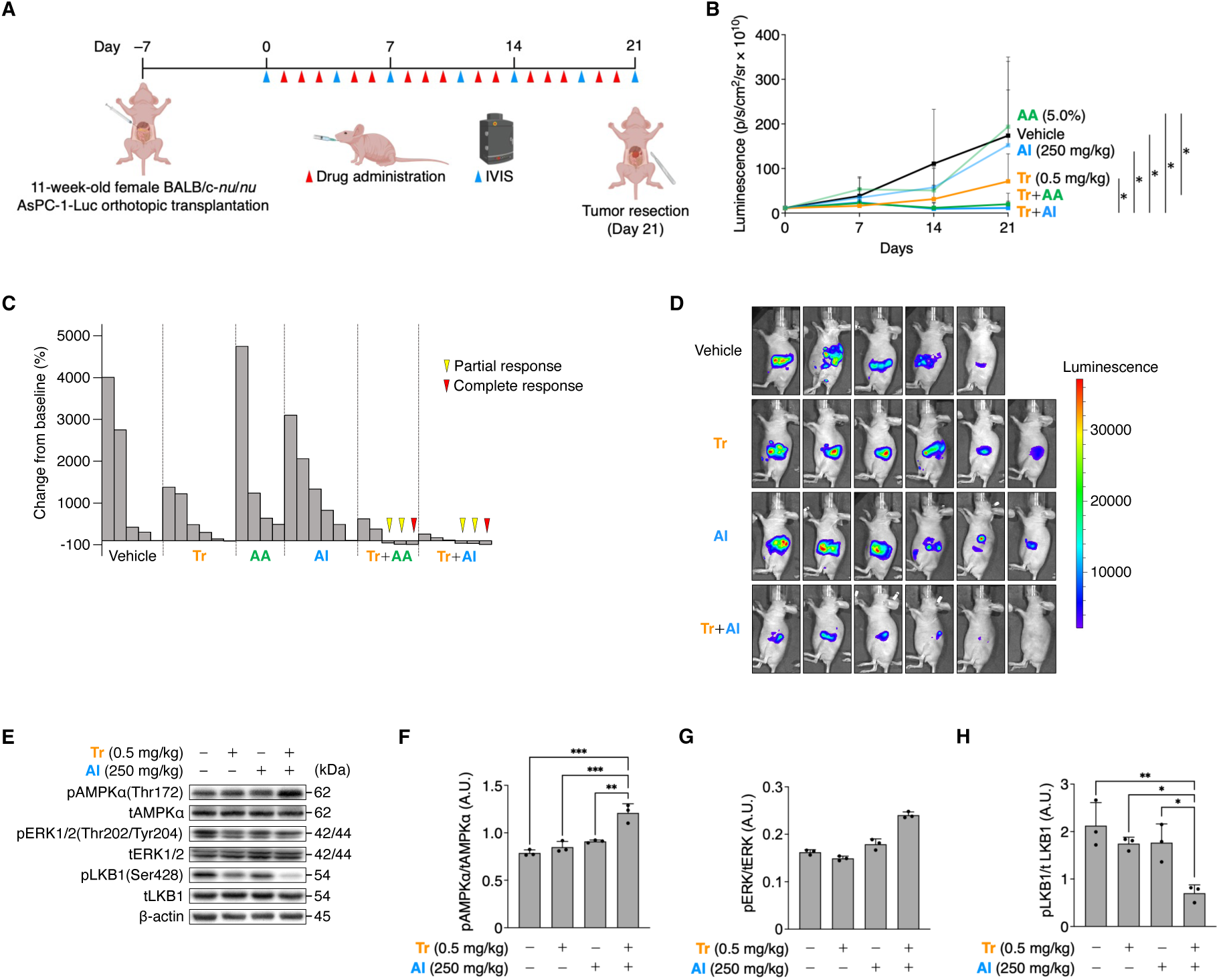
Co-targeting MAPK and AMPK signaling suppresses the growth of human PDAC xenografts. **A,** Schematic of the human PDAC xenograft experiment. Mice orthotopically transplanted with AsPC-1-Luc cells were randomised into six treatment groups (*n* = 6 per group) and treated orally five times per week for 3 weeks with vehicle, Tr (0.5 mg/kg), AA (5.0%), AI (250 mg/kg) or the indicated combinations (Tr+AA or Tr+AI). Tumor progression was monitored twice weekly using IVIS *in vivo* bioluminescence imaging. **B,** Combination treatment with Tr and AI results in a significantly greater reduction in tumor-associated bioluminescence (photons s^−1^ cm^−2^ sr^−1^) over the 21-day treatment period compared with the effect of Tr monotherapy. Tumor burden was quantified following luciferin administration. † indicates death during treatment (vehicle, *n* = 1; AA, *n* = 2; Tr+AA, *n* = 1). **C,** Co-targeting MEK and AMPK signaling induces tumor regression in a subset of mice, whereas monotherapies fail to elicit meaningful responses. Changes in tumor-associated bioluminescence at day 21 relative to baseline are categorized as partial or complete response. **D,** Representative bioluminescence images illustrating tumor progression from day 0 to day 21 across treatment groups. **E–H,** Tr+AI enhances AMPK activation and suppresses inhibitory phosphorylation of LKB1 in xenograft tumors. **E,** Immunoblotting of xenografts harvested at day 21 displaying phosphorylation of AMPKα, ERK and LKB1. β-actin served as a loading control. **F,** Quantification of the pAMPKα/tAMPKα ratio. **G,** Quantification of the pERK/tERK ratio. **H,** Quantification of the pLKB1/total LKB1 (tLKB1) ratio. Each dot represents an individual sample (**F–H**). Statistical significance was assessed by the Kruskal–Wallis test followed by the Steel–Dwass multiple comparisons test (**B**), and by one-way ANOVA followed by Tukey’s multiple comparisons test (**F–H**). \*\**P* < 0.01, \*\*\**P* < 0.001, \*\*\*\**P* < 0.0001. Data are presented as the mean ± SD. A.U., arbitrary units.

We therefore sought to identify pharmacologically tractable AMPK-activating compounds capable of recapitulating or exceeding the viability-rescuing effect of AA when combined with MEK inhibition. To this end, we screened small-molecule AMPK activators with established clinical safety profiles, either alone or in combination with 0.1 μM Tr. These compounds included bempedoic acid (BemA), metformin (Met), metformin hydrochloride (Met-HCl) and 5-aminoimidazole-4-carboxamide ribonucleoside (AICAR; AI). BemA has completed Phase III clinical trials and has been approved by the US Food and Drug Administration for the treatment of heterozygous familial hypercholesterolemia and atherosclerotic cardiovascular disease, whereas Met and Met-HCl have completed Phase IV clinical trials and are widely used for the treatment of T2DM. In addition, we tested AI, which has completed Phase III clinical trials for the prevention of ischemia–reperfusion injury in patients undergoing coronary artery bypass graft surgery. AI is a cell-permeable AMP analog that enters cells via adenosine transporters and is phosphorylated by adenosine kinase to 5-aminoimidazole-4-carboxamide ribonucleotide (ZMP), thereby activating AMPK in a manner analogous to AMP without directly altering intracellular ADP/ATP ratios or oxygen consumption, unlike many other AMPK activators.

After determining the maximum tolerated dose (MTD) of each compound in *Drosophila* (Figure S4A–D), we administered each AMPK activator in combination with Tr to *4-hit* flies. All four compounds exhibited significant synergistic effects with Tr, resulting in enhanced rescue of fly viability compared with any monotherapy (Figure 3B–E). Among these candidates, we selected AI for subsequent mechanistic analyses because its intracellular conversion to ZMP enables direct and specific activation of AMPK, facilitating more straightforward interpretation of AMPK-dependent signaling events. Consistent with the observed phenotypic synergy, immunoblotting revealed that combined Tr and AI treatment (Tr+AI) induced a stronger increase in AMPKα phosphorylation in *4-hit* larvae than either Tr or AI alone (Figure 3F, G). In parallel, Tr+AI robustly suppressed wing deformities in adult *4-hit* flies, phenocopying the corrective effects observed with the Tr+AA combination (Figure 3H, I).

Collectively, these findings demonstrate that clinically relevant AMPK activators can functionally substitute for microbiome-derived AA to synergise with MEK inhibition *in vivo*, reinforcing the concept that coordinated targeting of the MAPK and AMPK pathways represents an effective strategy for suppressing PDAC-associated oncogenic phenotypes.

### Co-targeting MAPK and AMPK pathways suppresses PDAC xenograft growth in mice

To evaluate whether the synergistic anti-tumor effect observed in *Drosophila* extend to a mammalian system, we next assessed the therapeutic efficacy of combined MEK and AMPK modulation in an orthotopic PDAC xenograft mouse model. Luciferase-labelled AsPC-1 cells (AsPC-1-Luc) were orthotopically implanted into the pancreatic tails of immunodeficient mice, enabling longitudinal monitoring of tumor burden by bioluminescence imaging. Following determination of dosing regimens based on MTD and PK analyses (Figure S5A–E), tumor-bearing mice were treated with AI or AA, either alone or in combination with Tr, and tumor growth was monitored over a three-week period (Figure 4A).

During treatment, one of six mice in the vehicle group and two of six mice in the AA monotherapy group died (Figure 4B). One mouse in the Tr+AA combination group also died, raising concerns regarding the tolerability of AA-based regimens in animals. At the experimental endpoint, mice treated with Tr+AI exhibited a significant reduction in tumor burden compared with vehicle-treated controls as well as with each monotherapy group (Figure 4B).

Consistent with these findings, tumor response analysis revealed no evidence of tumor regression in the AA or AI monotherapy groups, and only one of six mice treated with Tr alone exhibited tumor reduction (Figure 4C). Conversely, the Tr+AA combination significantly suppressed xenograft growth compared with AA monotherapy, with three of five mice exhibiting partial responses and one achieving a complete response (Figure 4B, C, Figure S5F). However, the occurrence of treatment-associated mortality underscores the limited therapeutic window of AA-based regimens. Notably, the Tr+AI combination induced partial responses in three of six mice and a complete response in one mouse without any treatment-related deaths (Figure 4C). Representative bioluminescence images confirmed that Tr+AI induced a marked and sustained reduction in tumor-associated luminescence relative to vehicle or monotherapy treatments (Figure 4D). Importantly, mice treated with Tr+AI did not exhibit significant body weight loss relative to vehicle-treated controls (Figure S5G), and all mice in the AI monotherapy and Tr+AI combination groups survived the treatment period, supporting the *in vivo* tolerability of this regimen.

To gain mechanistic insights into the anti-tumor effects of Tr+AI, we examined the activation status of key signaling components in xenograft tumors. Immunoblotting revealed robust phosphorylation of AMPKα in tumors from Tr+AI-treated mice (Figure 4E, F). Phosphorylation of ERK1/2 was largely comparable across treatment groups at the three-week time point (Figure 4G), suggesting that sustained MAPK pathway suppression may not be fully captured at this late stage. By contrast, phosphorylation of LKB1 at Ser428, an ERK-dependent inhibitory site, was markedly reduced in the Tr+AI group (Figure 4H), consistent with a mechanism whereby MEK inhibition relieves inhibitory regulation of the LKB1–AMPK axis, thereby facilitating AMPK activation.

Taken together, these findings demonstrate that co-targeting MAPK and AMPK signaling with Tr+AI effectively suppresses tumor growth in a PDAC xenograft model without notable toxicity. The associated enhancement of AMPK activation and reduction of inhibitory LKB1 phosphorylation provide mechanistic support for this combinatorial strategy and reinforce the therapeutic potential of coordinated modulation of the MAPK and AMPK pathways in PDAC.

### Co-targeting AMPK and MEK reduces markers of myofibroblastic differentiation of cancer-associated fibroblasts (myCAFs) and reshapes the stromal microenvironment

To investigate mechanisms underlying the anti-tumor effects of combined Tr and AI beyond direct tumor cell-intrinsic AMPK–MAPK signaling, we examined early stromal responses using an immunocompetent PDAC allograft model. KPC cells, derived from the autochthonous tumor of a KPC (*Pdx1-cre*;*LSL-Kras*^G12D^;*Trp53*^R172H/+^) mouse, were orthotopically transplanted into syngeneic C57BL/6J mice. Starting on day 6 post-transplantation, mice were treated by oral gavage with vehicle, Tr, AI or Tr+AI for four consecutive days. Tumors were harvested on day 10 for transcriptomic and histological analyses, enabling assessment of early treatment-induced changes before overt differences in the tumor burden emerged (Figure 5A).

**Figure 5.**
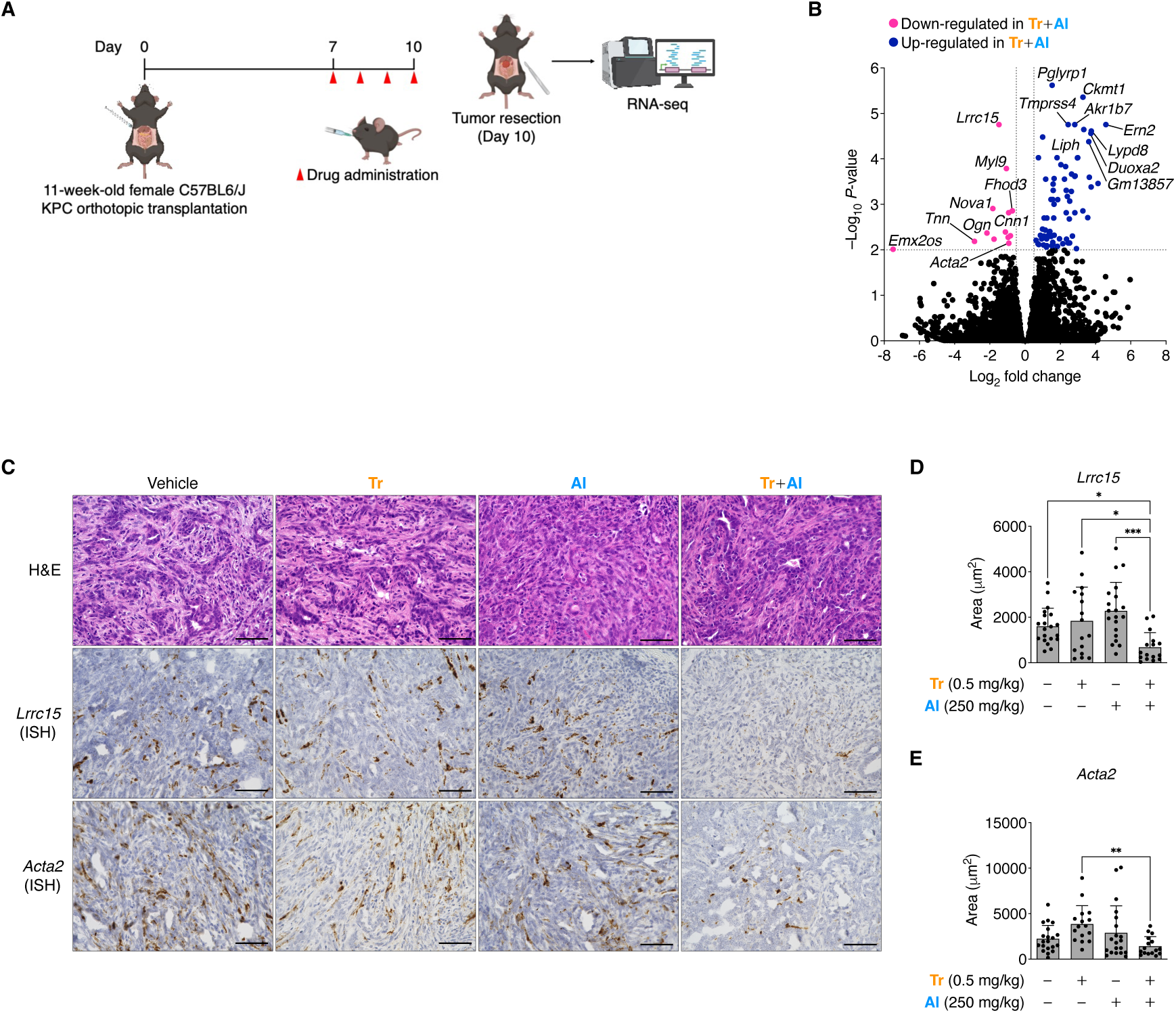
Tr+AI combination suppresses myCAFs and re-models the stroma in an immunocompetent allograft model. **A,** Schematic of the experimental design. KPC cells derived from genetically engineered PDAC mice were orthotopically transplanted into syngeneic C57BL/6J mice. Mice were randomised into four groups (*n* = 7 per group) and treated orally for 4 consecutive days, starting on day 7 post-transplantation, with vehicle, Tr (0.5 mg/kg), AI (250 mg/kg) or the Tr+AI combination. Tumors were harvested on day 10 for RNA-seq. **B,** Tr+AI combination selectively downregulates myCAF-associated genes, including *Lrrc15* and *Acta2*, compared with the effect of Tr monotherapy. Volcano plot presenting differentially expressed genes identified by RNA-seq analysis of Tr+AI versus Tr alone (|log_2_ fold change| ≥ 1; −log_10_ *P* ≥ 2). Genes down-regulated in the Tr+AI group are presented in magenta, and genes up-regulated in the Tr group are presented in navy. **C–E,** Tr+AI significantly suppresses expression of the myCAF markers *Lrrc15* and *Acta2*. **C,** Representative ISH images of *Lrrc15* and *Acta2* in serial sections of allograft tumors harvested on day 10. For each allograft, four randomly selected regions were imaged and evaluated. Scale bars, 100 μm. **D–E,** Quantification of the *Lrrc15*-positive (**D**) and *Acta2*-positive (**E**) areas. The Tr+AI group exhibits significantly reduced expression of both markers compared with the vehicle and monotherapy groups. Each dot represents a single microscopic field (**D–E**). Statistical significance was assessed using the Kruskal–Wallis test followed by Dunn’s multiple comparisons test (**D**), and by one-way ANOVA followed by Tukey’s multiple comparisons test (**E**). \*\**P* < 0.01, \*\*\**P* < 0.001. Data are presented as the mean ± SD.

RNA sequencing (RNA-seq) revealed significant downregulation of multiple markers associated with myCAFs, including *Lrrc15* and *Acta2*, in tumors from Tr+AI-treated mice compared with those treated with Tr alone (Figure 5B). myCAFs are characterized by high expression of LRRC15 and α-smooth muscle actin (encoded by *ACTA2*) and have been implicated in extracellular matrix (ECM) deposition and restriction of immune cell infiltration, contributing to the fibrotic and immunosuppressive tumor microenvironment typical of PDAC^37–39^. The selective reduction of myCAF-associated transcripts therefore suggests that Tr+AI may partially exert anti-tumor effects through suppression of pro-tumorigenic stromal fibroblast states linked to immune exclusion.

To spatially validate these transcriptomic findings, we performed *in situ* hybridisation (ISH) for *Lrrc15* and *Acta2*. Consistent with the RNA-seq results, ISH analysis demonstrated markedly reduced expression of both genes in tumors from the Tr+AI group compared with vehicle-, Tr- or AI-treated tumors (Figure 5C–E), supporting a selective inhibitory effect of the combination therapy on myCAF-associated features *in vivo*.

We next assessed broader stromal and vascular characteristics using histological, immunohistochemical and ISH analyses. Masson’s trichrome staining revealed increased collagen fiber deposition in Tr+AI-treated tumors, as well as in tumors treated with Tr or AI monotherapy, relative to vehicle-treated controls (Figure S6A, B). Notably, collagen deposition has been reported to exert contest-dependent effects in PDAC, including potential tumor-restraining roles^40^. Conversely, no significant differences were observed across treatment groups in *Col1a1* expression, CD31-positive vascular area or the proportion of moderately differentiated tumor regions (Figure S6A, C, D, E), indicating that Tr+AI does not induce broad stromal disruption or overt vascular re-modeling.

Analysis of additional fibroblast-associated markers revealed treatment-specific effects. Expression of *Islr*, a marker of undifferentiated fibroblasts^41^, was decreased in the Tr group but not in the Tr+AI group (Figure S6A, F), whereas expression of *Pi16,* another marker of immature fibroblasts^42^, was increased in the Tr group but remained unchanged in the combination group relative to vehicle (Figure S6A, G). These patterns suggest that although collagen deposition is enhanced across treatment conditions, Tr+AI preferentially influences CAF composition rather than globally altering fibroblast states.

Collectively, these findings indicate that co-targeting MEK and AMPK selectively modulates CAF composition, particularly by suppressing myCAF-associated features, while leaving vascular characteristics largely intact. Such selective stromal re-programming may contribute to the anti-tumor efficacy of Tr+AI by attenuating fibroblast-driven immune and physical barriers within the PDAC microenvironment.

### Dual inhibition of the TGF-β and MAPK pathways suppresses CAF-driven malignant phenotypes

To determine whether the anti-tumor effects of Tr+AI extend to direct suppression of CAF-mediated malignant traits, we focused on CAF differentiation and function using complementary cell-based assays. Differentiation of CAFs into myCAFs is tightly coupled to activation of the TGF-β signaling pathway^37^. Upon ligand binding, TGF-β receptors phosphorylate receptor-regulated Smads (R-Smads; Smad2 and Smad3), which form complexes with Smad4 and translocate into the nucleus to induce transcriptional programs associated with LRRC15^+^ myCAF differentiation^38^. Curiously, AMPK activation has been reported to inhibit TGF-β receptor–mediated phosphorylation of R-Smads, thereby attenuating downstream TGF-β signaling^43^.

Based on these observations, we hypothesised that concurrent MAPK inhibition and AMPK activation might co-operatively suppress TGF-β signaling programs in CAFs. To test this hypothesis, we first examined nuclear accumulation of phosphorylated Smad2/3 (pSmad2/3), a direct readout of TGF-β pathway activity. hPSC-5-mCherry cells, a human pancreatic stellate cell line labelled for lineage tracing, were stimulated with recombinant human TGF-β1 and analyzed by immunofluorescence (IF; Figure 6A). Compared with dimethyl sulfoxide (DMSO) controls, nuclear pSmad2/3 accumulation was significantly reduced in both Tr-and AI-treated cells (Figure 6B, C). Notably, Tr+AI induced a more pronounced reduction than either monotherapy, consistent with additive or cooperative suppression of TGF-β–Smad signaling (Figure 6B, C).

**Figure 6.**
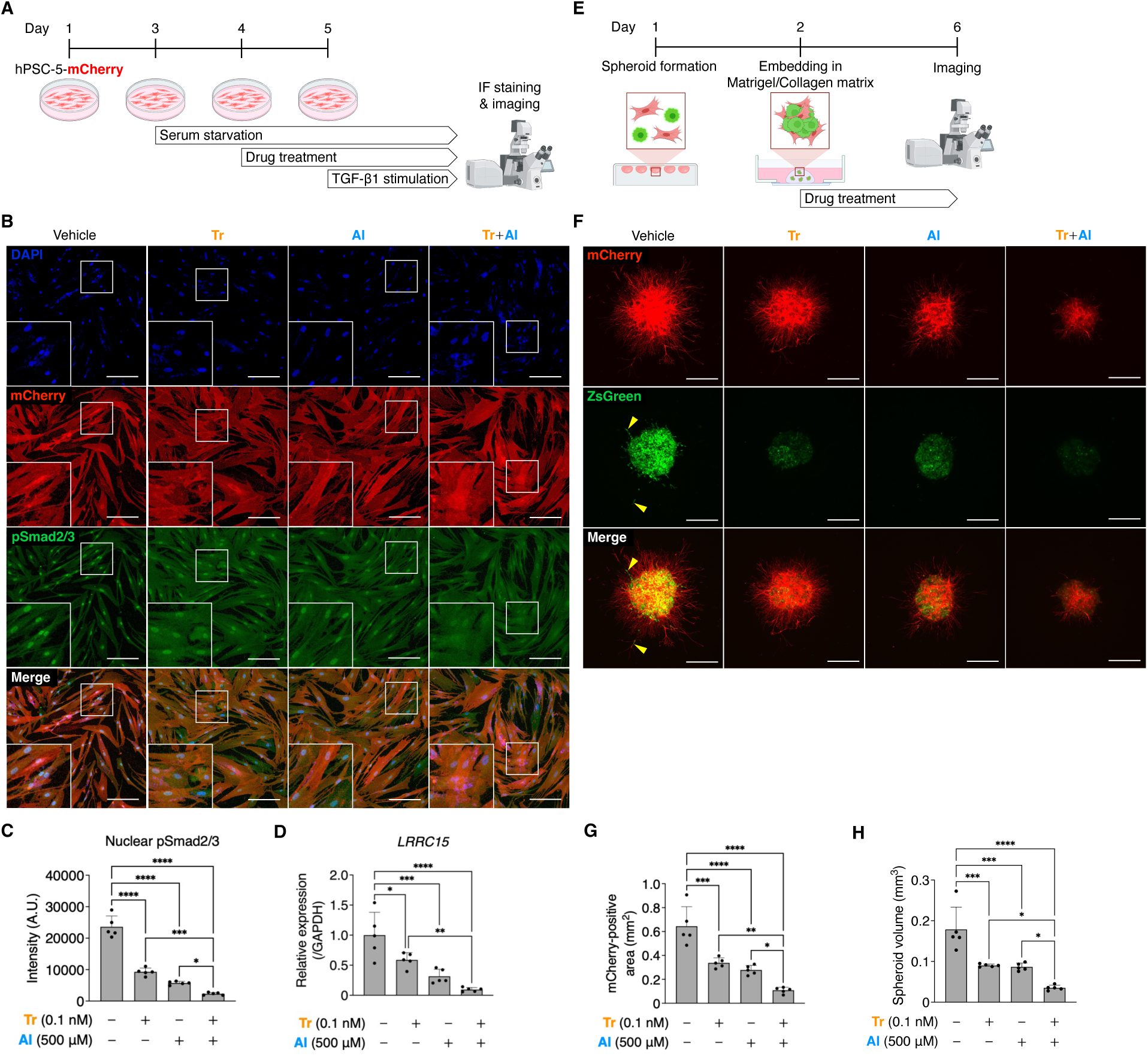
Dual modulation of MAPK and AMPK suppresses CAF-driven malignant phenotypes. **A,** Schematic of the IF experimental workflow. hPSC-5-mCherry CAFs were seeded on collagen I-coated glass-bottom dishes and cultured for 48 h. Cells were then serum-starved in low-serum medium (0.1% FBS) and treated for 24 h with DMSO, Tr (0.1 nM), AI (500 μM) or the Tr+AI combination. Recombinant TGF-β1 (10 ng/mL) was subsequently added for 60 min. Cells were fixed and subjected to IF staining for phosphorylated Smad2 and Smad3 (pSmad2/3) and DAPI. Confocal images were acquired using identical acquisition settings, and nuclear pSmad2/3 levels were quantified in mCherry-positive hPSC-5 CAFs. **B–C,** Tr+AI significantly suppresses nuclear translocation of pSmad2/3. Representative IF images of hPSC-5-mCherry CAFs stimulated with TGF-β1 following pre-treatment with DMSO, Tr, AI or Tr+AI. pSmad2/3 is presented in green, whereas mCherry-positive CAFs is presented in red and nuclei are presented in blue (**B**). Scale bars, 200 μm. Quantification of the mean nuclear pSmad2/3 fluorescence intensity is presented in (**C**). **D,** Tr+AI markedly suppresses TGF-β1–induced *LRRC15* expression in hPSC-5 CAFs, exceeding the effects of Tr or AI alone. Relative mRNA expression of *LRRC15*, a marker of myCAFs, was measured after TGF-β1 treatment (10 ng/mL, 48 h) under the indicated drug conditions and normalised to *GAPDH*. **E,** Schematic of the spheroid invasion assay. hPSC-5-mCherry CAFs (red) and AsPC-1-ZsGreen PDAC cells (green) were co-cultured and assembled into spheroids using the hanging-drop method. After 24 h, spheroids were embedded in a Matrigel/collagen I matrix and treated with DMSO, Tr (0.1 nM), AI (500 μM) or Tr+AI, followed by 4 days of culture with continuous drug exposure. **F–H,** Tr+AI exerts the strongest inhibitory effect on CAF-driven invasion and cancer cell proliferation. Representative maximum-intensity projections illustrate CAF-derived invasive strands and cancer cell infiltration into the matrix (**F**). Arrowheads indicate cancer cells infiltrating along CAF protrusions. Scale bars, 400 μm. Quantification of CAF invasion, measured as the mCherry-positive invasive area (**G**) and cancer cell proliferation, quantified as the ZsGreen-positive spheroid volume (**H**). The Tr+AI combination resulted in the greatest reduction in both parameters, indicating robust suppression of CAF-mediated invasion and cancer cell proliferation. Each dot represents an independent experiment (**C–D, G–H**). Boxed areas are presented at higher magnification in the insets. Statistical significance was assessed by one-way ANOVA followed by Tukey’s multiple comparisons test (**C–D, G–H**). \**P* < 0.05, \*\**P* < 0.01, \*\*\**P* < 0.001, \*\*\*\**P* < 0.0001. Data are presented as the mean ± SD. A.U., arbitrary units.

Next, we examined whether these signaling changes translated into altered myCAF marker expression. hPSC-5 cells were treated with TGF-β1 under the same drug conditions, and *LRRC15* expression was quantified by reverse transcription-qPCR (RT-qPCR). Both Tr and AI significantly reduced *LRRC15* expression compared with DMSO-treated controls, whereas Tr+AI further decreased *LRRC15* levels relative to either agent alone (Figure 6D). These results indicate that combined MAPK inhibition and AMPK activation more effectively attenuates TGF-β–driven myCAF gene expression than single-pathway modulation.

To determine the functional consequences of these molecular changes, we performed a three-dimensional spheroid invasion assay to assess CAF-driven invasion and cancer cell proliferation. hPSC-5-mCherry CAF and AsPC-1-ZsGreen PDAC cells were co-cultured as spheroids embedded in a Matrigel/collagen I matrix and treated with DMSO, Tr, AI or Tr+AI (Figure 6E). Under DMSO treatment, CAFs exhibited extensive radial invasion, generating prominent stromal tracks that facilitated infiltration of AsPC-1 cells (Figure 6F). Quantitative analyses confirmed that both CAF invasive area and cancer spheroid volume were greatest in the DMSO group (Figure 6G, H). Treatment with either Tr or AI alone significantly reduced CAF invasion and cancer cell proliferation, as evidenced by shorter invasive projections and reduced spheroid volumes (Figure 6F–H). AI treatment additionally reduced CAF expansion, consistent with attenuated TGF-β signaling. Strikingly, the Tr+AI combination exerted the strongest inhibitory effect: CAF-derived invasive projections were markedly diminished, and cancer cell proliferation was profoundly suppressed relative to all other treatment groups (Figure 6F–H). These finding indicate that concurrent MAPK inhibition and AMPK activation more effectively disrupts CAF-driven invasive and pro-tumorigenic behaviors than either intervention alone.

Finally, we assessed whether the stromal re-programming induced by Tr+AI represents a stable phenotypic change or a reversible state dependent on continued pathway modulation. To address this, hPSC-5-mCherry cells were pre-treated with DMSO, Tr, AI or Tr+AI for 48 h, followed by the addition of AsPC-1-ZsGreen cells and an additional 48 h of co-culture with or without continued drug treatment (Figure S7A). When drug treatment was maintained throughout the co-culture period, AsPC-1 proliferation was markedly suppressed by Tr or AI monotherapy and was most strongly inhibited by Tr+AI (Figure S7B, C). When drugs were withdrawn at the time of AsPC-1 seeding (Figure S7D, E), the suppressive effect on cancer cell proliferation was attenuated but not abolished, and significant differences among the treatment groups persisted. These findings indicate that transient CAF pre-conditioning is sufficient to partially restrain cancer cell proliferation, whereas sustained MAPK–AMPK co-targeting achieves more robust suppression in this co-culture context.

## DISCUSSION

PDAC remains among the most therapeutically refractory human malignancies, reflecting the convergence of oncogenic signaling rewiring, profound metabolic plasticity and a highly fibrotic, treatment-resistant tumor microenvironment. Although increasing evidence implicates the gut microbiome and its metabolites in shaping tumor behavior, how microbiome-derived metabolic cues are integrated with oncogenic signaling and stromal programs in PDAC remains poorly understood. The central conceptual advance of this study is the identification of reduced AA availability as a conserved metabolic constraint observed in PDAC that converges on suppression of AMPK signaling and can be therapeutically exploited through coordinated MEK–AMPK modulation. By integrating analyses of treatment-naïve patient samples with genetically tractable whole-animal models, we demonstrate a potential cross-scale mechanism linking microbial metabolism, tumor-intrinsic oncogenic signaling and stromal plasticity in PDAC.

SCFAs, including AA, PA and BA, are recognised for their immunomodulatory and metabolic effects, yet their roles in cancer are highly context dependent. Although SCFAs can promote anti-inflammatory immune responses and tumor suppression in some settings, they may also be co-opted by metabolically stressed tumors as alternative carbon sources^44^. Our observation that both the relative abundance of putative AA-producing taxa and fecal or systemic AA levels are consistently reduced in PDAC patients and a in a neoplastic animal model supports the view that diminished AA availability represents a disease-associated metabolic feature linked to PDAC-related oncogenic programs, rather than an artifact of experimental manipulation. Importantly, our human cohort consisted exclusively of treatment-naïve patients with PDAC, capturing endogenous microbiome and metabolite alterations inherently associated with disease state. When integrated with our *Drosophila* data demonstrating that oncogenic genotypes directly constrain colonization by AA-producing bacteria, these findings suggest a model in which tumor-intrinsic oncogenic signaling reshapes the microbiome–metabolite landscape, rather than microbiome changes acting as a primary initiating event.

Beyond cancer cell-intrinsic effects, our study reveals a previously underappreciated role for MEK–AMPK co-targeting in stromal re-programming. CAFs are central regulators of PDAC progression, ECM re-modeling, immune exclusion and therapeutic resistance^45,46^. Among CAF subtypes, myCAFs, characterized by high *LRRC15* and *ACTA2* expression, promote a fibrotic and immunosuppressive tumor microenvironment and correlate with poor responses to immunotherapy across multiple cancer types, including PDAC^38,39^. In an immunocompetent PDAC allograft model, combined MEK–AMPK modulation selectively suppressed myCAF markers without broadly ablating stromal compartments including the vasculature. Mechanistically, this effect was linked to attenuation of TGF-β–Smad2/3 signaling in CAFs, reduced *LRRC15* expression and diminished CAF invasiveness. These findings are particularly relevant given prior studies that indiscriminate fibroblasts depletion can paradoxically accelerate PDAC progression^47^, underscoring the importance of stromal normalization rather than stromal elimination. In this context, our findings raise the possibility that MEK–AMPK co-targeting alleviates hostile tumor microenvironmental features characteristic of PDAC by selectively decreasing pro-tumorigenic myCAF states, a hypothesis that warrants direct evaluation in future studies.

Several limitations of this work should be acknowledged. Although our integrated analyses support a functional link among microbiome-associated AA availability, AMPK signaling and PDAC progression, establishing definitive causal relationships will require validation in longitudinal human cohorts and gnotobiotic or genetically controlled mammalian models. In addition, whether AMPK dependence varies across molecular PDAC subtypes or treatment histories remains to be determined.

In summary, our work delineates a PDAC-associated AA reduction and links this metabolic alteration to impaired AMPK signaling within oncogenic contexts. Rather than positioning AA itself as a therapeutic agent, our findings identify AMPK as an actionable signaling hub at the interface of metabolism, oncogenic pathways and stromal regulation. More broadly, this work underscores how integrating microbiome-informed metabolic states with pathway-oriented intervention can inform rational treatment strategies for PDAC and other desmoplastic, therapeutically refractory malignancies.

## RESOURCE AVAILABILITY

### Lead contact

Further information and requests for resources should be directed to and will be fulfilled by the lead contact, Masahiro Sonoshita.

### Materials availability

Further information and requests for resources and reagents should be directed to and will be fulfilled by the lead contact, Masahiro Sonoshita. All unique/stable reagents generated in this study are available from the lead contact upon reasonable request and with a completed materials transfer agreement (MTA).

### Data and code availability

Datasets containing human participant data are not publicly available due to ethical restrictions regarding patient confidentiality, but are available from the corresponding author upon reasonable request. The 16S rRNA sequencing data and RNA-seq data generated during the present study are also available from the corresponding author upon reasonable request. Source data for all Figures are provided with the paper.

## ACKNOWLEDGMENTS

We thank all study participants and the medical staff at Hokkaido University Hospital for their invaluable cooperation and support. We are also grateful to the members of the Sonoshita Laboratory for insightful discussions and continuous contributions to this work. We also thank Dr. Fumiaki Obata and members of his laboratory for their thoughtful discussions and constructive input.

This research was funded by the Japan Society for the Promotion of Science (JSPS) through grants 21K14762 and 24K17810 (awarded to R.Y.), 22H03541 (to S.F.), 20H03524, 23H02759, and PJS00420230001 (to M.S.), the Japan Science and Technology Agency (JST) through grants JPMJCR2333 (to M.S.) and R03W01 (to R.Y.), Suzuken Memorial Foundation, Extramural Collaborative Research Grant of Cancer Research Institute at Kanazawa University, Takeda Science Foundation, Suhara Memorial Foundation, Ono Medical Research Foundation, the NOVARTIS Foundation (Japan) for the Promotion of Science, Daiichi-Sankyo ‘Habataku’ Support Program for the Next Generation of Researchers, the Japanese Cancer Association and Kobayashi Foundation for Cancer Research (to R.Y.), Japan Agency for Medical Research and Development-Core Research for Evolutionary Science and Technology (AMED-CREST) through grant JP23gm1010009, JST-Exploratory Research for Advanced Technology (JST-ERATO) through grant JPMJER1902, and the Food Science Institute Foundation (to S.F.), Takeda Science Foundation, G-7 Scholarship Foundation, the Junior Scientist Promotion and Photo-Excitonix projects at Hokkaido University, and the Joint Research Program of the Institute for Genetic Medicine, Hokkaido University (to M.S.).

## AUTHOR CONTRIBUTIONS

R.Y. and M.S. conceived the study and supervised the project. R.Y., J.F., T.Ki., T.O., S.S., S.K., K.H., Y.K., K.K., M.K., S.T., M.W., T.A., T.N., N.S. and S.Hi. conceived and conducted the clinical research, obtained patient consent and collected human fecal samples and clinical data. T.S. and S.F. performed SCFA quantification of human fecal samples by GC/MS. R.Y. performed microbiome and metabolome data analyses. A.V.T. performed compositional analysis of the microbiome data, and R.E.L. provided funding for A.V.T. T.O., K.C.H., Y.H., T.Mi. and Y.M. constructed and performed pathological evaluation of patient TMA. R.Y., Y.S., T.H. and S.Ha. carried out all fly experiments. R.Y., S.J. and K.M. quantified short-chain fatty acids in fly samples by LC-MS/MS. R.Y., Y.S., T.Ki., T.O., T.H., S.Ha., R.S., C.K. and T.Mo. performed xenograft and allograft mouse experiments. R.Y. performed downstream data analysis of RNA-seq data. Y.U. and A.E. conducted histopathological analyses of allograft tumors. R.Y. and T.Ka. performed spheroid invasion assays and two-dimensional co-culture experiments of cancer cells and CAFs. R.Y. wrote the manuscript with input from all authors.

## DECLARATION OF GENERATIVE AI AND AI-ASSISTED TECHNOLOGIES

During the preparation of this work, the authors used Gemini (Google) and ChatGPT (OpenAI) solely to refine the English phrasing and improve the clarity of the manuscript. After using these tools, the authors reviewed and edited the content as needed and take full responsibility for the content of the publication.

## STAR METHODS

### KEY RESOURCES TABLE

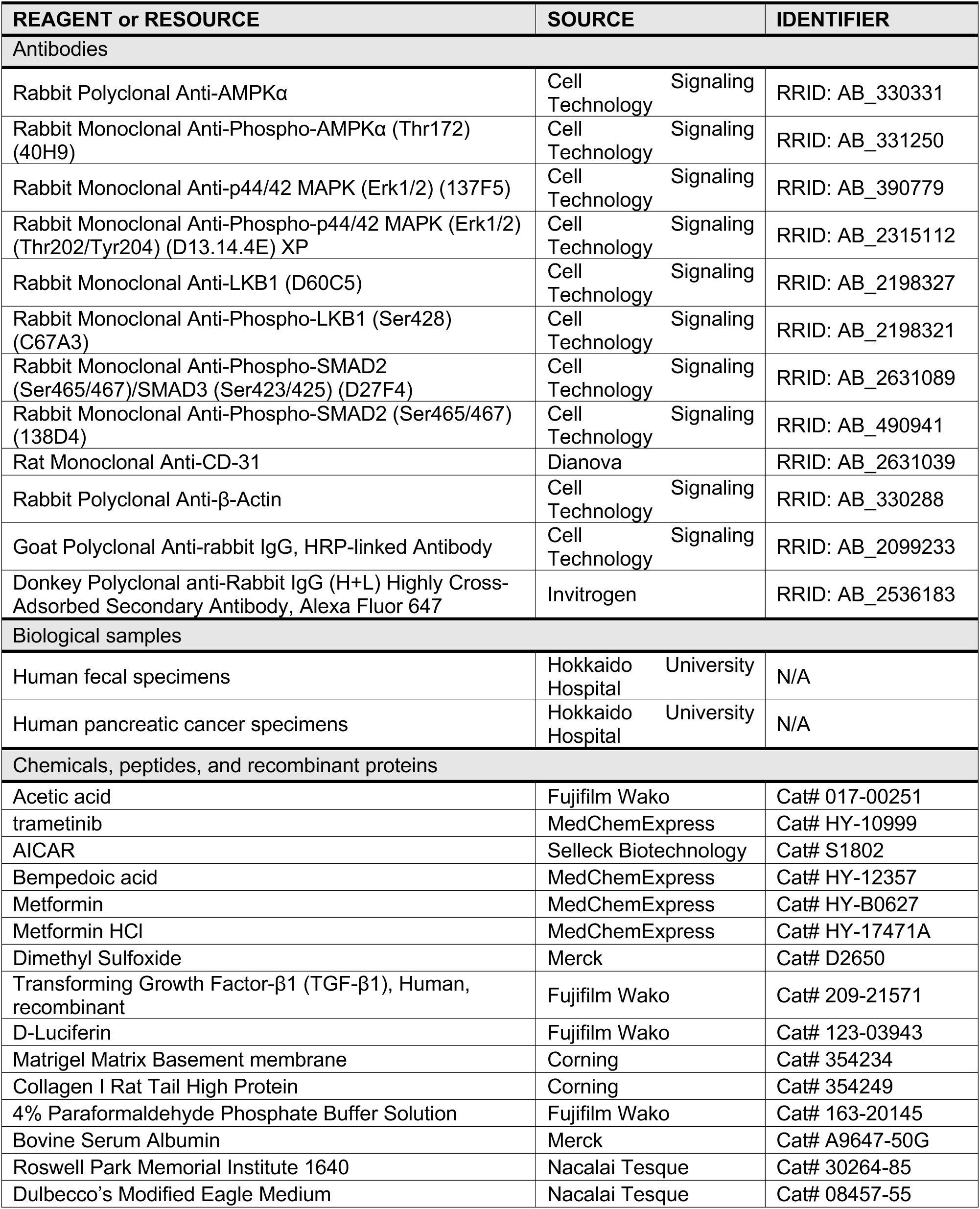
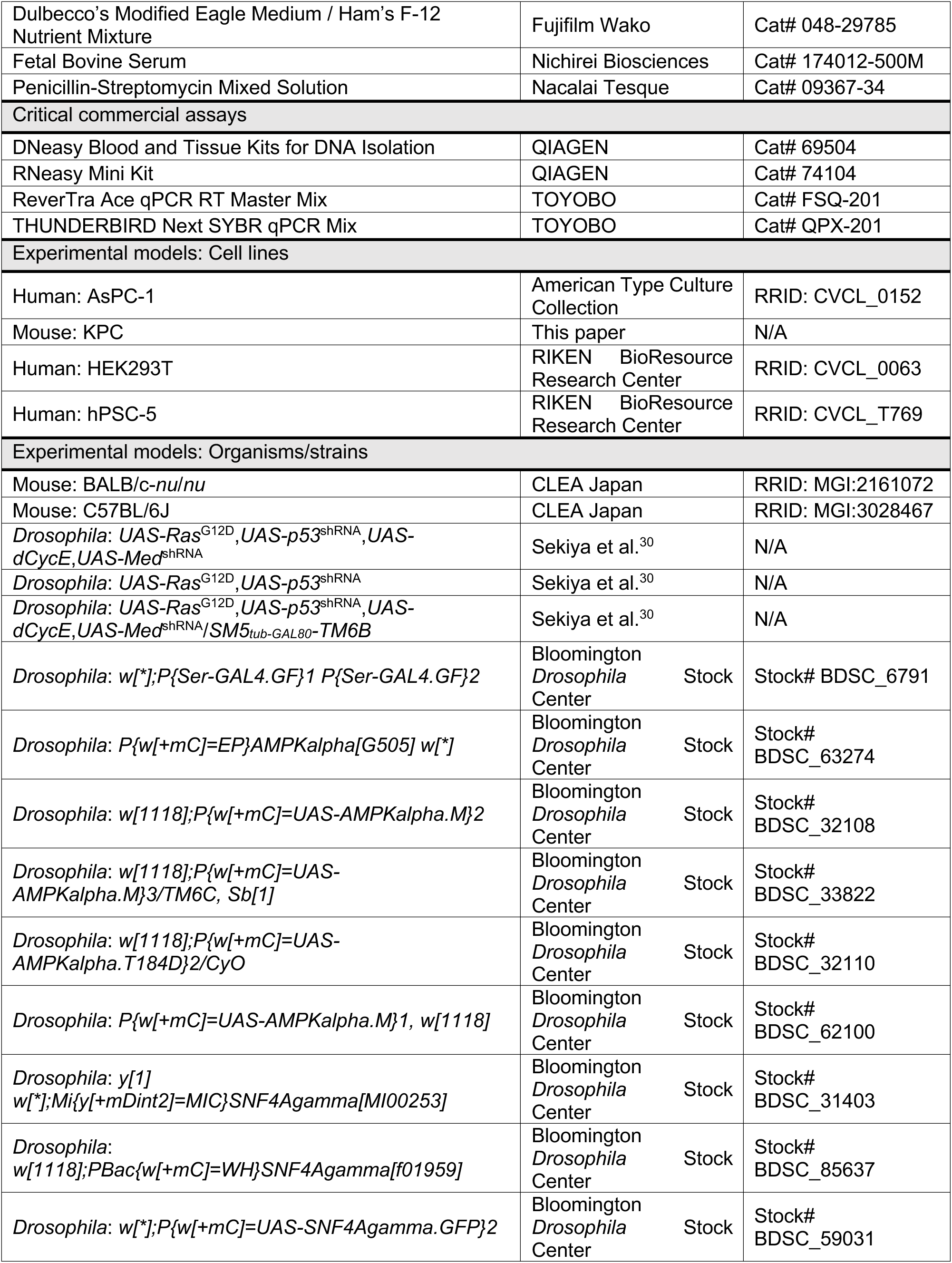
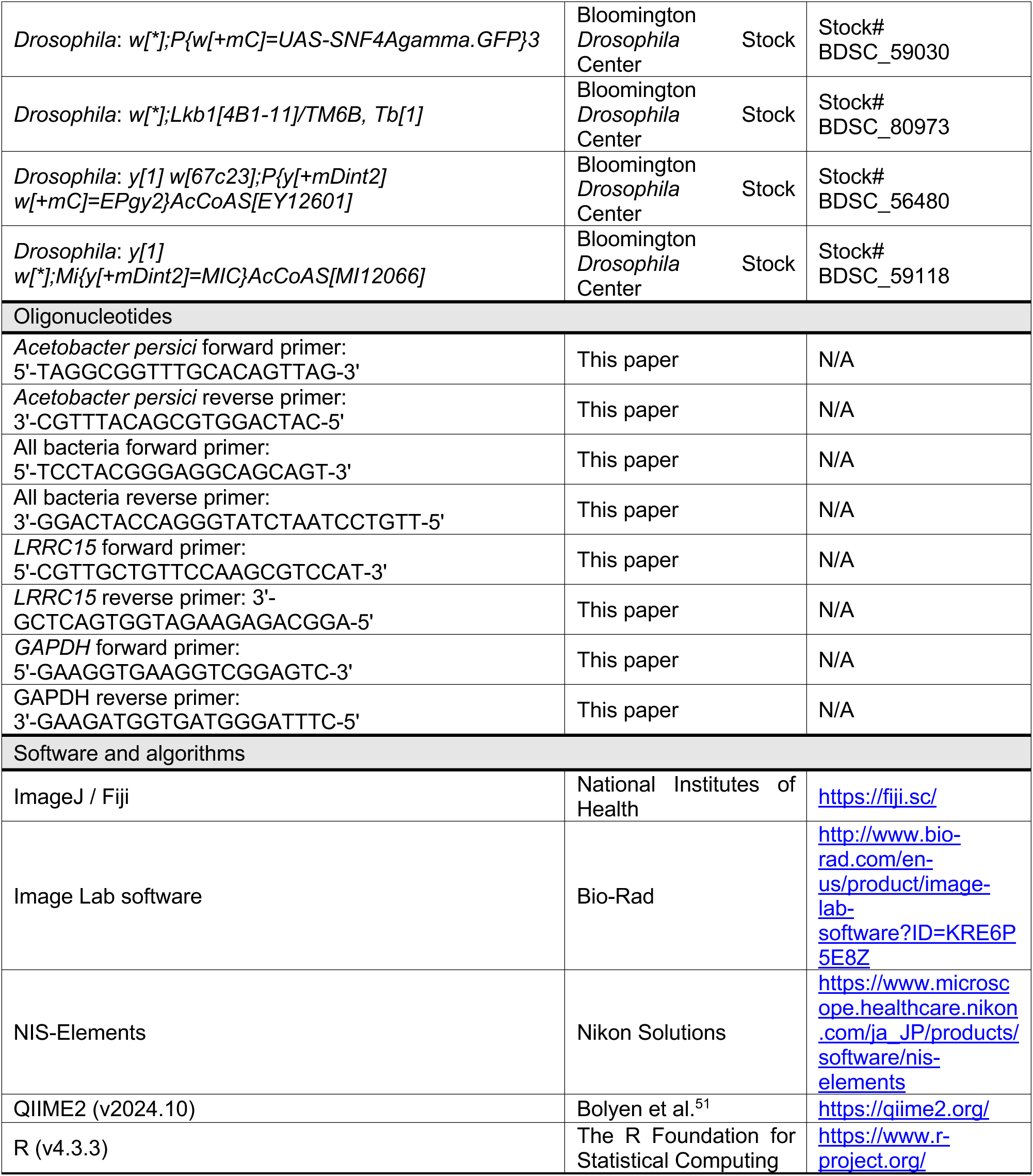

### EXPERIMENTAL MODEL AND STUDY PARTICIPANT DETAILS

#### Human study participants and fecal samples

In total, 44 patients diagnosed with PDAC were consecutively recruited at the Departments of Gastroenterology and Hepatology and Gastroenterological Surgery II at Hokkaido University Hospital (Sapporo, Japan). Patients were treated either as outpatients or inpatients (PDAC group). Fecal samples were successfully obtained from 33 of the 44 participants (collection rate: 75.0%). In parallel, 42 HCs provided written informed consent, and fecal samples were collected from all volunteers (collection rate: 100%). The eligibility criteria for the PDAC group included age ≥20 years, pathological confirmation of adenocarcinoma and no antibiotic exposure within 3 months prior to sample collection. HCs were required to be ≥20 years old, have no history of serious gastrointestinal disease, report no gastrointestinal symptoms at the time of enrolment and have no antibiotic exposure within 3 months. At outpatient consultation or hospital admission, participants received two fecal collection kits (one containing guanidine thiocyanate [GuSCN] solution [Technosuruga Laboratory, Shizuoka, Japan] and one lacking GuSCN), along with detailed instructions, a cooler bag and refrigerant packs. Participants collected fresh fecal samples, stored them in the cooler bag with pre-frozen refrigerant and mailed them to the hospital on the following day. Upon receipt, samples intended for DNA extraction were mixed with GuSCN solution and stored at 4°C until processing, whereas samples without GuSCN were immediately stored at −80°C for gas chromatography/mass spectrometry (GC/MS).

All participants were of Japanese descent, and their demographic characteristics, including sex and gender, are summarized in Table S3. All procedures involving human participants in the study involving fecal sample collection were conducted in accordance with the Declaration of Helsinki and approved by the Ethics Committee of Hokkaido University Hospital (approval number: 020–0355). Written informed consent was obtained from all participants.

#### Human PDAC tissue specimens for TMA construction

A TMA was constructed using tumor tissue specimens from 153 patients with PDAC (Figure 3J). To minimise confounding effects of postoperative treatment, 67 patients who received adjuvant chemotherapy were excluded, and an additional eight patients were excluded because of insufficient evaluable tumor tissue, resulting in 78 patients in the final analysis. Immunohistochemistry (IHC) was performed using antibodies against AMPKα (RRID: AB_330331; 1:100; Cell Signaling Technology, Danvers, MA, USA) and phospho-AMPKα (Thr172) (RRID, AB_331250; 1:100; Cell Signaling Technology). TMA cores were independently evaluated by two blinded investigators: T.O. (a board-certified pathologist of the Japanese Society of Pathology) and R.Y. For each patient, the proportion of AMPKα-positive or pAMPKα-positive cancer cells was quantified across the entire core. Patients were categorized as marker-positive or marker-negative, and associations with overall survival, disease-free survival and other clinicopathologic characteristics were analyzed.

All participants were of Japanese descent, and their demographic characteristics, including sex and gender, are summarized in Table S3. TMA analysis was approved by the Ethics Committee of Hokkaido University Hospital (approval number: 019–0154) and conducted in accordance with institutional ethical guidelines. Tumor specimens were collected under a broad consent for future research, and for deceased patients, surrogate consent was obtained in accordance with institutional policy.

#### Generation of *4-hit* flies modeling the PDAC genotype

The generation of the *4-hit Drosophila* line was previously described^30^. In brief, cDNA encoding constitutively active *Drosophila Ras*^G12D^ and a short hairpin RNA (shRNA) targeting *p53* were cloned into an expression vector containing two *upstream activation sequence* (*UAS*) sites^48^. The resulting plasmid was injected into *y1w67c23*;*P{CaryP}attP2* embryos to establish the *UAS*-*Ras*^G12D^,*UAS*-*p53*^shRNA^ line. In parallel, HA-tagged *Drosophila Cyclin E* (*dCycE*) and an shRNA targeting *Medea* (*Med*; the *Drosophila* orthologue of *SMAD4*) were cloned into an empty *UAS* vector and injected into *PBac {yellow[*+*]-attP-9A} VK00027* embryos to generate the *UAS-dCycE*,*UAS-Med*^shRNA^ line. These two strains were subsequently crossed, and recombinant progeny were selected to generate the quadruple transgenic *UAS-Ras*^G12D^, *UAS-p53*^shRNA^,*UAS-dCycE*,*UAS-Med*^shRNA^ (*UAS-4-hit*) line. To drive tissue-specific expression of the four transgenes, *UAS-4-hit* flies were crossed with the *Serrate* (*Ser*)*-gal4* driver line, yielding *Ser-gal4*;*UAS-Ras*^G12D^,*UAS-p53*^shRNA^,*UAS-dCycE*,*UAS-Med*^shRNA^ (*4-hit*) flies.

All *Drosophila* studies were conducted in accordance with protocols approved by Hokkaido University Safety Committee on Genetic Recombination Experiments (approval numbers: 2019–007 and 2022–029).

#### *Drosophila* medium preparation

Brewer’s yeast (MP Biomedicals, Solon, OH, USA), yeast extract (Sigma-Aldrich), Bacto Casitone (Thermo Fisher Scientific), agar, sucrose, glucose, MgCl_2_, CaCl_2_, PA and mould inhibitor (10% methyl-4-hydroxybenzoate in 95% ethanol; Fujifilm Wako) were dissolved in reverse osmosis-purified water. The mixture was brought to a boil and allowed to cool prior to use.

#### Mice

Female BALB/c-*nu/nu* mice (11-week-old; CLEA) and female C57BL/6J mice (11-week-old; CLEA) were housed under specific pathogen-free conditions with a 12-h/12-h light/dark cycle. For xenograft assays, mice were euthanized when humane endpoints approved by the Institutional Animal Care and Use Committee were reached. These endpoints included signs of severe distress; development of ascites, tumor metastasis or gastrointestinal bleeding (e.g. blood in stool); weight loss of ≥15%; and tumor diameter exceeding 2 cm.

All animal experiments were conducted in accordance with protocols approved by the Hokkaido University Safety Committee on Genetic Recombination Experiments (approval numbers: 2019–007 and 2022–029) and the Hokkaido University Animal Research Committee (approval numbers: 19–0121, 2022–0117 and 2022–0118).

#### Cell lines

The human pancreatic cancer cell line AsPC-1 (derived from a male patient) and the murine pancreatic cancer cell line KPC (derived from a male mouse) were maintained in Roswell Park Memorial Institute (RPMI) 1640 Medium (Nacalai Tesque, Kyoto, Japan). HEK293T cells (derived from a male patient) were cultured in Dulbecco’s Modified Eagle Medium (DMEM; Nacalai Tesque), while the human pancreatic stellate cell line hPSC-5 (sex unknown) was cultured in Dulbecco’s Modified Eagle Medium / Ham’s F-12 Nutrient Mixture (DMEM/F-12; Fujifilm Wako). All culture media were supplemented with 10% heat-inactivated fetal bovine serum (FBS; Nichirei Biosciences) and 100 U/mL penicillin-streptomycin (Nacalai Tesque). All cells were incubated at 37°C in a humidified atmosphere with 5% CO_2_. We verified the absence of *Mycoplasma* contamination through routine testing and authenticated all cell lines via annual short tandem repeat profiling. To ensure cellular integrity, we visually assessed cell morphology before each experiment and during cryopreservation.

### METHOD DETAILS

#### 16S rRNA gene sequencing and bioinformatics analysis

Bacterial DNA was extracted from human fecal samples and from *Drosophila* midgut and hindgut tissues as previously described^49^. The V3–V4 region of the bacterial 16S rRNA gene was amplified and sequenced using the Illumina MiSeq platform (Illumina, San Diego, CA, USA) following established protocols^50^. Bioinformatics, including quality filtering, denoising and downstream analysis, was performed using QIIME2^51^. Paired-end FASTQ reads were demultiplexed, and amplicon sequence variants were inferred using the DADA2 tool. Taxonomic classification was performed across five hierarchical ranks (phylum, class, order, family, genus) using the SILVA 132 reference database at 99% sequence similarity threshold.

Differential abundance analysis at the genus level was performed using the analysis of composition of microbiomes with bias correction (ANCOM-BC)^52^ plugin in QIIME2, following the standard tutorial workflow (https://docs.qiime2.org/2024.10/tutorials/moving-pictures/). ANCOM-BC accounts for compositionality and corrects for biases due to unequal sampling depths; *q*-values were calculated using the Benjamini–Hochberg false discovery rate correction to adjust for multiple comparisons. Taxa with *q* of <0.05 were considered significantly differentially abundant between HC and PDAC patient samples, and separately between control and *4-hit* flies. To further account for potential confounding factors, we applied multi-variable association with linear models (MaAsLin2)^53^ to taxa identified as differentially abundant by ANCOM-BC in human samples. Furthermore, two models were tested: one adjusted for age and sex; and another adjusted for age, sex, BMI and type 2 diabetes mellitus (T2DM) status.

To complement the taxon-specific analysis, we also performed a compositional-aware investigation to capture broader structural differences in the microbiome between groups. To achieve this, the taxonomic composition tables (read counts) were filtered by prevalence (prevalence ≥ 50% at abundance ≥ 0.05%) and aggregated at the genus level. Moreover, taxa annotated as ‘unclassified’ were discarded. Further analysis was conducted similarly to that described previously^54^. Briefly, normalization was conducted using *clr*-transformation. General association of microbiome composition and sample group was tested using a PERMANOVA test (9999 permutations) based on compositional Aitchison beta-diversity metric. We applied the Nearest Balance (NB) method^8^ to assess microbiome compositional shifts between HC and PDAC patient samples, as well as between the microbiomes of control and *4-hit* flies.

#### GC/MS

Nine organic acids—FA, AA, PA, IBA, BA, IVA, VA, LA, and SA—were quantified in fecal samples using an Agilent 5977B GC/MS system (Agilent Technologies, Santa Clara, CA, USA), following established protocols^55^. Briefly, fecal samples were lyophilised for at least 18 h and homogenised with 3.0-mm zirconia beads at 1500 × *g* for 10 min, and 10 mg of the processed fecal material was used for each analysis.

#### LC-MS/MS

SCFAs in *Drosophila* larvae were quantified by LC-MS/MS using a previously described method^56^. Briefly, 50 mg of larval tissue were washed with phosphate-buffered saline (PBS; Fujifilm Wako, Osaka, Japan), accurately weighed, flash-frozen in liquid nitrogen and stored at −80°C until analysis. For extraction, 250 μL of 99.5% ethanol (Fujifilm Wako) were added to each frozen sample, followed by homogenisation using a bead crusher (TAITEC, Koshigaya, Japan) at 2500 rpm for 4 min. Homogenates were centrifuged at 12,000 rpm for 10 min at 4°C and 50 μL of the resulting supernatant were collected. To this aliquot, we added 50 μL each of 2 mg/L ethyl 2-ethylbutyrate (internal standard; Sigma-Aldrich, St. Louis, MO, USA), 50 mM 3-nitrophenylhydrazine, 50 mM 1-ethyl-3-(3-dimethylaminopropyl)carbodiimide (Sigma-Aldrich) and 7.5% pyridine (Sigma-Aldrich), followed by 50 μL of 75% aqueous methanol (Sigma-Aldrich). Samples were vortexed, incubated at 25°C for 30 min in the dark and then diluted 5-fold with 75% aqueous methanol containing 0.5% FA (Sigma-Aldrich). SCFAs were analyzed on a Dionex Ultimate 3000 LC system coupled to a TSQ Quantum Access Max triple quadrupole mass spectrometer (Thermo Fisher Scientific, Waltham, MA, USA), as previously described^56^.

#### qPCR

Third-instar larvae were collected from plastic vials and sequentially washed for 30 s each in ice-cold 70% ethanol (Fujifilm Wako), ice-cold PBS (Fujifilm Wako), RNA*later* Stabilization Solution and ice-cold PBS. Midgut and hindgut tissues were dissected from 10 larvae, pooled into a single sample in PBS, and stored at −80°C until DNA extraction. Genomic DNA was extracted using a DNeasy Blood and Tissue Kit (QIAGEN) following the manufacturer’s instructions. Quantification of AP was performed by qPCR as described previously^33^. Reaction mixtures were prepared using THUNDERBIRD SYBR qPCR Mix (Toyobo, Osaka, Japan) and run on a StepOnePlus real-time PCR system (Thermo Fisher Scientific). Universal 16S ribosomal DNA primers served as an internal control. The qPCR cycling conditions were 95°C for 1 min, followed by 40 cycles of 95°C for 15 s and 60°C for 45 s. Relative AP abundance was calculated using the 2^−ΔΔCt^ method.

#### Immunoblotting

For total protein extraction from *Drosophila*, five third-instar larvae were pooled. Mouse tissues were minced into approximately 1-mm³ tissue fragments prior to lysis. Protein concentrations were determined using the Bradford assay. Equal amounts of total protein were resolved on 10% SDS-PAGE gels and transferred to polyvinylidene difluoride (PVDF) membranes (MilliporeSigma, Burlington, MA, USA). Membranes were blocked with Blocking One or Blocking One P (Nacalai Tesque) for 2 h at room temperature and incubated with primary antibodies overnight at 4°C. After washing, membranes were probed with goat anti-rabbit IgG secondary antibodies. Immunoreactive bands were imaged using the ChemiDoc XRS+ system and quantified with Image Lab software (Bio-Rad Laboratories, Hercules, CA, USA). Relative signal intensities were quantified using β-actin as a loading control and normalized to the total protein levels of the corresponding targets.

#### *Drosophila* chemical testing

AA (Fujifilm Wako) was dissolved in double-distilled water. All other compounds, namely Tr, AI, BemA, Met and Met-HCl, were dissolved in DMSO (Sigma-Aldrich) to prepare stock solutions. Fly food was supplemented with vehicle and/or chemicals at a final DMSO concentration of 0.1% and dispensed into plastic vials (Thermo Fisher Scientific).

Before viability assays, the MTD of each compound was determined by assessing the survival of non-transgenic control (*white*^−^) flies fed each chemical. For experimental crosses, virgin *UAS-4-hit* females were crossed with *Ser*-*gal4* males to generate *4-hit* offspring. These flies were reared at 22°C for assays involving AA monotherapy, Tr+AA combination treatment and wing area measurements or at 23.5°C for assays testing individual AMPK activators alone or in combination with Tr. Percent viability was calculated as the number of adults enclosed divided by the total number of pupae. Adult wings were dissected and wing area was quantified using ImageJ software (US National Institutes of Health, MD, USA).

#### *Drosophila* genetic screening

Fly stocks carrying kinase gene mutations were obtained from Bloomington *Drosophila* Stock Center. To ensure genetic consistency, balancer chromosomes were standardised before screening. Stocks balanced with *FM6*, *FM7a*, *FM7c* or *FM7i* were crossed with the *FM7c-Tb* (*Tb*)*-RFP* balancer line. *CyO*-, *SM5*- and *SM6a*-balanced stocks were re-balanced using *CyO*-*Tb*-*RFP*, whereas *TM3*-, *TM6C*- and *MKRS*-balanced stocks were re-balanced using the *TM6B* balancer carrying the visible *Tb* marker.

For genetic screening of kinase genes located on the X chromosome, *4-hit* males were crossed with either control (*w*^−^) or kinase-mutant females. To assess the heterozygosity of kinase genes on the autosomes, *4-hit* females were crossed with control or kinase-mutant males harboring mutations on the second, third or fourth chromosome. For gain-of-function studies, *4-hit* females were crossed with males carrying *UAS*-*gene*^OE^, generating *4-hit* progeny in which gene expression was induced specifically in GAL4-expressing cells.

All progeny, irrespective of kinase heterozygosity, knockdown or over-expression, were cultured on standard fly food at 27°C for 11 days until adulthood. Percent viability was determined as described for the chemical screening assays. *Drosophila* orthologues of human genes were predicted using *Drosophila* RNAi Screening Center Integrative Ortholog Predictive Tool (https://www.flyrnai.org/cgi-bin/DRSC_orthologs.pl).

#### PDAC mouse models

A PDAC xenograft model was established as previously described^31^. In brief, 1 × 10^6^ luciferase-transduced AsPC-1 cells were orthotopically injected into the pancreatic tails of anesthetized female BALB/c-*nu*/*nu* mice (11-week-old; CLEA, Tokyo, Japan). Mice were randomised into six treatment groups (*n* = 6 per group) with comparable baseline tumor size, as determined by bioluminescence imaging using the IVIS Spectrum Imaging System (PerkinElmer, Waltham, MA, USA). Animals received oral treatment with vehicle, Tr (0.5 mg/kg, MedChemExpress, Monmouth Junction, NJ, USA), AA (5.0%), AI (250 mg/kg, Selleck Chemicals, Houston, TX, USA) or the indicated combinations (Tr+AA or Tr+AI) five times per week for 3 weeks. Tumor growth was monitored by bioluminescence imaging twice weekly and quantified using Living Image software (v4.2; PerkinElmer).

For syngeneic allograft assays, 1 × 10^6^ KPC cells, which were derived from the autochthonous tumor of a KPC (*Pdx1-cre*;*LSL-Kras*^G12D^;*Trp53*^R172H/+^) mouse, were injected into the pancreatic tails of anesthetized female C57BL/6J mice (11-week-old; CLEA Japan, Osaka, Japan). Mice were randomised into four groups (*n* = 7 per group) and treated orally once daily for 4 consecutive days, starting 7 days after transplantation, with vehicle, Tr (0.5 mg/kg), AI (250 mg/kg) or Tr+AI.

#### RNA-seq

Tumor tissues were harvested from the pancreas of allograft mice immediately after completion of the 4-day treatment regimen and immersed in RNA*later* Stabilization Solution (Thermo Fisher Scientific) to preserve RNA integrity. Total RNA was extracted using RNeasy Mini Kit (QIAGEN, Hilden, Germany), and RNA quality and integrity were assessed using NanoDrop spectrophotometer (Thermo Fisher Scientific) and Agilent Bioanalyzer (Agilent Technologies).

Poly(A)^+^ mRNA was purified using the Poly(A) mRNA Magnetic Isolation Module (New England Biolabs, Ipswich, MA, USA), and sequencing libraries were prepared with the NEBNext Ultra II Directional RNA Library Prep Kit (New England Biolabs). Libraries were sequenced on a NovaSeq X Plus platform (Illumina) to generate paired-end reads. Raw sequencing reads were quality-checked using FastQC (v0.11.7) and trimmed with Trimmomatic (v0.38). Clean reads were aligned to the mouse reference genome assembly GRCm38 (mm10) using HISAT2 (v2.1.0). Gene-level read counts were generated using featureCounts (v1.6.3), and transcript abundance was calculated as transcripts per million. Differential gene expression analysis was conducted using DESeq2 (v1.24.0) with relative log expression normalization. Genes with |log_2_ fold change| > 1 and adjusted *P* < 0.05 (Benjamini–Hochberg correction) were considered differentially expressed. Gene Ontology enrichment analysis was performed using GOATOOLS (v1.1.6), with multiple testing corrections applied using the Benjamini–Hochberg method.

#### Histological analysis of mouse tumors

Formalin-fixed, paraffin-embedded mouse tumor tissues were sectioned at 4 µm and subjected to RNA ISH, IHC or Masson’s trichrome staining.

For RNA ISH, assays were conducted using the RNAscope 2.5 HD Reagent Kit–Brown (Advanced Cell Diagnostics, Newark, CA, USA). Sections were deparaffinised, treated with hydrogen peroxide for 10 min at room temperature and boiled in target retrieval solution for 15 min using a steam warmer or for 3 min using a pressure cooker (SR-MP300; Panasonic, Osaka, Japan). After incubation with Protease Plus (Advanced Cell Diagnostics) for 30 min at 40°C, sections were hybridised with target probes for 2 h at 40°C in a HybEZ II Hybridisation System (Advanced Cell Diagnostics). Signal amplification was performed using AMP 1–6 reagents, and signal detection was conducted using 3,3′-diaminobenzidine (DAB), followed by hematoxylin counterstaining. The stained sections were dehydrated, cleared and mounted using Entellan mounting media (Merck, Darmstadt, Germany). The following RNAscope probes were used: *Mm-Islr* (NM_012043.4, region: 763–1690, Cat No. 450041), *Mm-Lrrc15* (NM_028973.2, region: 514–1649, Cat No. 467831), *Mm-Acta2* (NM_007392.3, region: 41–1749, Cat No. 319531), *Mm-Pi16* (NM_023734.3, region: 731–1846, Cat No. 451311) and *Mm-Col1a1* (NM_007742.3, region: 1686–4669, Cat No. 319371).

For IHC, sections were deparaffinised, rehydrated and subjected to antigen retrieval in Epitope Retrieval Solution pH 6 (Leica Biosystems, Wetzlar, Germany) for 30 min using a pressure cooker. After blocking with 2.5% normal goat or horse serum (Vector Laboratories, Newark, CA, USA), slides were incubated overnight at 4°C with anti-CD31 antibody (RRID: AB_2631039; 1:50; Dianova, Hamburg, Germany) diluted in Immuno Shot Reagent 1 (Cosmo Bio, Tokyo, Japan). Endogenous peroxidase activity was quenched using 3% H_2_O_2_ (Fujifilm Wako), and signals were visualised using the ImmPRESS HRP Goat Anti-Rat IgG Polymer Detection Kit (Vector Laboratories) and ImmPACT DAB EqV Substrate, Peroxidase (Vector Laboratories). The slide sections were counterstained with hematoxylin.

Masson’s trichrome staining was performed using standard protocols. In brief, deparaffinised and rehydrated tissue sections were immersed in a 1:1 (v/v) mixture of 10% trichloroacetic acid (Fujifilm Wako) and 10% potassium dichromate (Fujifilm Wako) for 30 min, followed by rinsing under running tap water. The sections were then stained with Weigert’s iron hematoxylin for 2–3 min. After washing, sections were incubated in a 1:1 (v/v) mixture of 2.5% phosphomolybdic acid (Tokyo Chemical Industry, Tokyo, Japan) and 2.5% phosphotungstic acid (Alfa Aesar, Heysham, UK) for 45 s and stained with 0.75% Orange G (Fujifilm Wako). The sections were washed with 1% AA and further stained in a solution of Ponceau-S (Sigma-Aldrich), fuchsin acid (Fujifilm Wako) and azophloxin (Sigma-Aldrich) for 30 min. After another wash with 1% AA, the sections were placed in 2.5% phosphotungstic acid for 10 min. Finally, the washed sections were stained in aniline blue (Kishida Chemical, Osaka, Japan) for 10 min and quickly washed again with 1% AA.

Brightfield microscopy images were acquired using a BZ-X710 microscope (Keyence, Osaka, Japan) with a ×40 objective lens. For each sample, DAB-positive areas (for ISH and IHC) and blue-stained fibrotic areas (for Masson’s trichrome) were quantified in four randomly selected fields using BZ-X Analyser software (Keyence).

#### Generation of AsPC-1 cells stably expressing exogenous markers

AsPC-1-Luc cells were generated as previously described^33^. We generated AsPC-1-ZsGreen cells using a lentiviral vector (pLVSIN-EF1α-IRES-ZsGreen1; Takara Bio, Shiga, Japan) encoding luciferase cDNA and ZsGreen. The plasmid was co-transfected with the Lentiviral High Titer Packaging Mix into HEK293T cells using ViaFect (Promega, Madison, WI, USA) following the manufacturer’s protocol. At 48 h post-transfection, viral supernatants were collected, filtered and used to infect AsPC-1 cells in the presence of polybrene (4.0 µg/mL). Infected cells were trypsinized and seeded at clonal density (1.0 × 10^3^ cells per 100-mm dish). ZsGreen-positive colonies were identified by fluorescence microscopy, isolated and expanded, and luciferase expression was subsequently confirmed.

#### Generation of hPSC-5 cells stably expressing exogenous markers

hPSC-5 cells (RCB3588) were transduced with mCherry using a lentiviral transduction system. In brief, HEK293T cells were seeded in 100-mm dishes and cultured to 80%–90% confluence on the day of transfection. Lentiviral plasmids pCSII-mCherry-Hyg, pVSVG, pREV and pRRE were co-transfected into HEK293T cells using ViaFect (Promega) following the manufacturer’s protocol. After 24 h, the medium was replaced with DMEM containing 10% FBS and 2 mM caffeine (Thermo Fisher Scientific). Viral supernatants were harvested 48 h post-transfection, filtered through a 0.45-µm PVDF membrane and mixed 1:1 with fresh culture medium. The filtered supernatant was applied to hPSC-5 cells for transduction, and transduced cells were selected with hygromycin B (300 µg/mL; InvivoGen, San Diego, CA, USA) until hygromycin B-resistant colonies emerged. Stable mCherry expression was verified using fluorescence microscopy, and positive populations were expanded.

#### IF staining

hPSC-5-mCherry cells were cultured in 24-well glass-bottom dishes (MatTek, Ashland, MA, USA) pre-coated with collagen I (100 µg/mL; Corning, Corning, NY, USA). Cells were seeded at a density of 5.0 × 10^4^ cells per well in DMEM/Ham’s F-12 medium (Fujifilm Wako). After 48 h, cells were treated with DMSO, Tr (0.1 nM), AI (500 µM) or Tr+AI. Twenty-four hours later, the medium was replaced with low-serum medium [0.1% FBS (Nichirei Biosciences)] containing the corresponding treatments, and cells were serum-starved for an additional 24 h.

Recombinant human TGF-β1 (10 ng/mL; Fujifilm Wako) was then added while maintaining drug exposure. After 60 min of TGF-β1 stimulation, cells were fixed with 4% paraformaldehyde (Fujifilm Wako) for 15 min at room temperature and washed three times with PBS. Cells were permeabilised and blocked in PBS containing 5% BSA with 0.3% Triton X-100 1 h at room temperature, followed by overnight incubation at 4°C with a primary antibody against pSmad2/3. After three washes with PBS, fluorophore-conjugated secondary antibodies and DAPI (Dojindo Laboratories, Kumamoto, Japan) were applied for 60 min at room temperature in the dark.

Fluorescence images were acquired using a Nikon A1R HD25 confocal microscope (Nikon, Tokyo, Japan) using identical exposure conditions to avoid signal saturation. For quantification, five randomly selected fields per well were imaged from five independent wells per condition. Nuclear pSmad2/3 levels were quantified using ImageJ by measuring the mean fluorescence intensity within DAPI-defined nuclei of mCherry-positive hPSC-5 fibroblasts, and values were compared across treatment groups.

#### Spheroid invasion assay

Spheroid invasion assays were performed as previously described^57^ with minor modifications. AsPC-1-ZsGreen and hPSC-5-mCherry cells were suspended in DMEM/Ham’s F-12 and mixed with sterile 1.2% methylcellulose solution (cP 4000; Sigma-Aldrich) prepared in the same medium at a 1:1:1 ratio. Approximately 5.0 × 10^3^ cells of each cell type were seeded in 20-µL hanging drops on the inner surface of 100-mm dish lids (50 droplets per lid) and incubated for 24 h at 37°C in a humidified 5% CO_2_ incubator to allow spheroid formation. The following day, a gel mixture consisting of collagen I (4 mg/mL) and Matrigel (2 mg/mL; Corning) was prepared by neutralised with 5× Collagen stock solution (100 mM HEPES pH7.5, 2% NaHCO_3_, 50 mg/mL αMEM). Spheroids were collected by gently tapping the lids and rinsing them with 1 mL of medium, transferred to 1.5-mL tubes and allowed to settle by gravity for 30 s. After removing the supernatant, spheroids were resuspended in 200 µL of the gel mixture and dispensed as 40-µL droplets into 24-well glass-bottom dishes. The dishes were inverted every 45 s until gelation (approximately 15 min), followed by an additional 30-min incubation at 37°C. Complete medium containing DMSO, Tr, AI or Tr+AI was then gently overlaid onto the gels and maintained throughout the culture period, with a single medium change on day 2.

Spheroids were cultured for 4 days and imaged on day 6 using a Nikon A1R HD25 confocal microscope equipped with a ×10 objective, acquiring z-stacks at 0.75-µm intervals. Image stacks were processed into maximum-intensity projections using NIS-Elements software (Nikon). For quantitative analysis, CAF invasion was assessed by measuring the mCherry-positive invasive area of hPSC-5 cells from maximum-intensity projections using ImageJ. Cancer cell proliferation was quantified by measuring the ZsGreen-positive spheroid volume of AsPC-1 cells, calculated as *V* = 4/3π[(*d*_max_+*d*_min_)/4]^3^, where *d*_max_ and *d*_min_ represent the major and minor spheroid diameters, respectively. For each condition, five wells were analyzed, with five randomly selected fields per well.

#### Two-dimensional co-culture of hPSC-5-mCherry and AsPC-1-ZsGreen

hPSC-5-mCherry cells were cultured in 24-well glass-bottom dishes pre-coated with collagen I (100 µg/mL). Cells were seeded at a density of 5.0 × 10^4^ cells/mL in DMEM/Ham’s F-12 medium. After 48 h, cultures were treated with DMSO, Tr (0.1 nM), AI (500 µM) or Tr+AI. Following 48 h of compound treatment, AsPC-1-ZsGreen cells were added to the same wells at a density of 5.0 × 10^4^ cells/mL in the same medium, with or without continued drug exposure. After an additional 48 h of co-culture, fluorescence images were acquired using a confocal microscope (A1R HD25) under identical exposure conditions to avoid signal saturation. For each condition, five wells were analyzed, with five randomly selected microscopic fields per well. The number of AsPC-1-ZsGreen cells was quantified using ImageJ.

### QUANTIFICATION AND STATISTICAL ANALYSIS

All statistical analyses were performed using R v4.3.3 (The R Foundation for Statistical Computing, Vienna, Austria) and GraphPad Prism v10.2.2 (GraphPad Software, Boston, MA, USA). The statistical tests used, exact sample sizes (*n*), and the definitions of *n* (e.g., individual human subjects, animals, pooled fly samples, independent experiments, or microscopic fields) are detailed in the corresponding figure legends. Data are presented as mean ± SD, mean ± SEM, or median with IQR, as indicated in the legends. For comparisons between two groups, a two-sided Student’s *t*-test was used. For multiple-group comparisons, one-way ANOVA followed by Tukey’s or Dunnett’s post hoc test was applied when assumptions for parametric testing were met; otherwise, non-parametric tests were used. Specifically, the Kruskal–Wallis test followed by Steel–Dwass, Dunn’s, or Steel’s multiple comparisons test was applied as indicated. Survival curves were estimated using the Kaplan–Meier method and compared using the log-rank test. Microbial differential abundance was assessed using ANCOM-BC and MaAsLin2, with MaAsLin2 models adjusted for age, sex, BMI, and T2DM status. Overall microbial community structure was evaluated by PERMANOVA based on weighted UniFrac distances, and compositional shifts were assessed using the NB method. The normality of the data was assessed to determine the appropriate statistical tests. All tests were two-tailed, and *P* < 0.05 was considered statistically significant.

## REFERENCES

1. Siegel, R.L., Giaquinto, A.N., and Jemal, A. (2024). Cancer statistics, 2024. CA Cancer J. Clin. 74, 12–49.

2. Park, E.M., Chelvanambi, M., Bhutiani, N., Kroemer, G., Zitvogel, L., and Wargo, J.A. (2022). Targeting the gut and tumor microbiota in cancer. Nat. Med. 28, 690–703.

3. Dickson, I. (2018). Microbiome promotes pancreatic cancer. Nat. Rev. Gastroenterol. Hepatol. 15, 328.

4. Tucker, O.N., Dannenberg, A.J., Yang, E.K., and J., F.I.T. (2004). Bile acids induce cyclooxygenase-2 expression in human pancreatic cancer cell lines. Carcinogenesis 25, 419–423.

5. Panebianco, C., Villani, A., Pisati, F., Orsenigo, F., Ulaszewska, M., Latiano, T.P., Potenza, A., Andolfo, A., Terracciano, F., Tripodo, C., et al. (2022). Butyrate, a postbiotic of intestinal bacteria, affects pancreatic cancer and gemcitabine response in in vitro and in vivo models. Biomed. Pharmacother. 151, 113163.

6. Temel, H.Y., Kaymak, Ö., Kaplan, S., Bahcivanci, B., Gkoutos, G.V., and Acharjee, A. (2023). Role of microbiota and microbiota-derived short-chain fatty acids in PDAC. Cancer Med. 12, 5661–5675.

7. Kuever, J. (2014). The Family Desulfovibrionaceae. In The Prokaryotes (Springer Berlin Heidelberg), pp. 107–133.

8. Odintsova, V.E., Klimenko, N.S., and Tyakht, A.V. (2022). Approximation of a microbiome composition shift by a change in a single balance between two groups of taxa. mSystems 7, e0015522.

9. Furusawa, Y., Obata, Y., Fukuda, S., Endo, T.A., Nakato, G., Takahashi, D., Nakanishi, Y., Uetake, C., Kato, K., Kato, T., et al. (2013). Commensal microbe-derived butyrate induces the differentiation of colonic regulatory T cells. Nature 504, 446–450.

10. Luu, M., Riester, Z., Baldrich, A., Reichardt, N., Yuille, S., Busetti, A., Klein, M., Wempe, A., Leister, H., Raifer, H., et al. (2021). Microbial short-chain fatty acids modulate CD8+ T cell responses and improve adoptive immunotherapy for cancer. Nat. Commun. 12, 4077.

11. Koh, A., De Vadder, F., Kovatcheva-Datchary, P., and Backhed, F. (2016). From Dietary Fiber to Host Physiology: Short-Chain Fatty Acids as Key Bacterial Metabolites. Cell 165, 1332–1345.

12. Han, K.-I., Kim, J.-S., Eom, M.K., Lee, K.C., Suh, M.K., Kim, H.S., Park, S.-H., Lee, J.H., Kang, S.W., Park, J.-E., et al. (2021). Collinsella acetigenes sp. nov., an Anaerobic Actinobacterium Isolated from Human Feces, and Emended Description of the Genus Collinsella and Collinsella aerofaciens. Curr. Microbiol. 78, 3667–3673.

13. Lan, Q., Liufu, S., Chen, B., Wang, K., Chen, W., Xiao, L., Liu, X., Yi, L., Liu, J., Xu, X., et al. (2025). Gut-resident Phascolarctobacterium succinatutens decreases fat accumulation via MYC-driven epigenetic regulation of arginine biosynthesis. NPJ Biofilms Microbiomes 11, 150.

14. Schug, Z.T., Vande Voorde, J., and Gottlieb, E. (2016). The metabolic fate of acetate in cancer. Nat. Rev. Cancer 16, 708–717.

15. Moore, F., Weekes, J., and Hardie, D.G. (1991). Evidence that AMP triggers phosphorylation as well as direct allosteric activation of rat liver AMP-activated protein kinase. A sensitive mechanism to protect the cell against ATP depletion: A sensitive mechanism to protect the cell against ATP depletion. Eur. J. Biochem. 199, 691–697.

16. Shen, C.H., Yuan, P., Perez-Lorenzo, R., Zhang, Y., Lee, S.X., Ou, Y., Asara, J.M., Cantley, L.C., and Zheng, B. (2013). Phosphorylation of BRAF by AMPK impairs BRAF-KSR1 association and cell proliferation. Mol. Cell 52, 161–172.

17. Jones, R.G., Plas, D.R., Kubek, S., Buzzai, M., Mu, J., Xu, Y., Birnbaum, M.J., and Thompson, C.B. (2005). AMP-Activated Protein Kinase Induces a p53-Dependent Metabolic Checkpoint. Mol. Cell 18, 283–293.

18. Inoki, K., Zhu, T., and Guan, K.-L. (2003). TSC2 mediates cellular energy response to control cell growth and survival. Cell 115, 577–590.

19. Faubert, B., Boily, G., Izreig, S., Griss, T., Samborska, B., Dong, Z., Dupuy, F., Chambers, C., Fuerth, B.J., Viollet, B., et al. (2013). AMPK is a negative regulator of the Warburg effect and suppresses tumor growth in vivo. Cell Metab. 17, 113–124.

20. Esteve-Puig, R., Canals, F., Colomé, N., Merlino, G., and Recio, J.Á. (2009). Uncoupling of the LKB1-AMPKα Energy Sensor Pathway by Growth Factors and Oncogenic BRAFV600E. PLoS One 4, e4771.

21. Zheng, B., Jeong, J.H., Asara, J.M., Yuan, Y.-Y., Granter, S.R., Chin, L., and Cantley, L.C. (2009). Oncogenic B-RAF negatively regulates the tumor suppressor LKB1 to promote melanoma cell proliferation. Mol. Cell 33, 237–247.

22. Sueda, T., Sakai, D., Kawamoto, K., Konno, M., Nishida, N., Koseki, J., Colvin, H., Takahashi, H., Haraguchi, N., Nishimura, J., et al. (2016). BRAFV600E inhibition stimulates AMP-activated protein kinase-mediated autophagy in colorectal cancer cells. Sci. Rep. 6, 18949.

23. Dar, A.C., Das, T.K., Shokat, K.M., and Cagan, R.L. (2012). Chemical genetic discovery of targets and anti-targets for cancer polypharmacology. Nature 486, 80–84.

24. Sonoshita, M., and Cagan, R.L. (2017). Modeling human cancers in Drosophila. In Current topics in developmental biology (Elsevier), pp. 287–309.

25. Sonoshita, M., Scopton, A.P., Ung, P.M.U., Murray, M.A., Silber, L., Maldonado, A.Y., Real, A., Schlessinger, A., Cagan, R.L., and Dar, A.C. (2018). A whole-animal platform to advance a clinical kinase inhibitor into new disease space. Nat. Chem. Biol. 14, 291–298.

26. Yamamura, R., Ooshio, T., and Sonoshita, M. (2021). Tiny Drosophila makes giant strides in cancer research. Cancer Sci. 112, 505–514.

27. Jiang, H., Kimura, T., Hai, H., Yamamura, R., and Sonoshita, M. (2022). Drosophila as a toolkit to tackle cancer and its metabolism. Front. Oncol. 12, 982751.

28. Hirabayashi, S., Baranski, T.J., and Cagan, R.L. (2013). Transformed Drosophila cells evade diet-mediated insulin resistance through wingless signaling. Cell 154, 664–675.

29. Yamamoto, M., Ohsawa, S., Kunimasa, K., and Igaki, T. (2017). The ligand Sas and its receptor PTP10D drive tumour-suppressive cell competition. Nature 542, 246–250.

30. Sekiya, S., Fukuda, J., Yamamura, R., Ooshio, T., Satoh, Y., Kosuge, S., Sato, R., Hatanaka, K.C., Hatanaka, Y., Mitsuhashi, T., et al. (2023). Drosophila Screening Identifies Dual Inhibition of MEK and AURKB as an Effective Therapy for Pancreatic Ductal Adenocarcinoma. Cancer Res. 83, 2704–2715.

31. Fukuda, J., Kosuge, S., Satoh, Y., Sekiya, S., Yamamura, R., Ooshio, T., Hirata, T., Sato, R., Hatanaka, K.C., Mitsuhashi, T., et al. (2024). Concurrent targeting of GSK3 and MEK as a therapeutic strategy to treat pancreatic ductal adenocarcinoma. Cancer Sci. 10.1111/cas.16100.

32. Jiang, H., Satoh, Y., Yamamura, R., Ooshio, T., Luo, Y., Hai, H., Otsuka, T., Hata, S., Sato, R., Hirata, T., et al. (2025). Inhibition of NAD-GPx4 axis and MEK triggers ferroptosis to suppress pancreatic ductal adenocarcinoma. Mol. Ther. 33, 4618–4635.

33. Qian, Z.R., Rubinson, D.A., Nowak, J.A., Morales-Oyarvide, V., Dunne, R.F., Kozak, M.M., Welch, M.W., Brais, L.K., Da Silva, A., Li, T., et al. (2018). Association of Alterations in Main Driver Genes With Outcomes of Patients With Resected Pancreatic Ductal Adenocarcinoma. JAMA Oncol 4, e173420.

34. Crotti, E., Rizzi, A., Chouaia, B., Ricci, I., Favia, G., Alma, A., Sacchi, L., Bourtzis, K., Mandrioli, M., Cherif, A., et al. (2010). Acetic acid bacteria, newly emerging symbionts of insects. Appl. Environ. Microbiol. 76, 6963–6970.

35. Nakao, K.-I., Ro, A., and Kibayashi, K. (2014). Evaluation of the morphological changes of gastric mucosa induced by a low concentration of acetic acid using a rat model. J. Forensic Leg. Med. 22, 99–106.

36. Subramanian, S., Du, C., and Tan, X.-D. (2022). Can rodent model of acetic acid-induced colitis be used to study the pathogenesis of colitis-associated intestinal fibrosis? J. Invest. Surg. 35, 223–224.

37. Biffi, G., Oni, T.E., Spielman, B., Hao, Y., Elyada, E., Park, Y., Preall, J., and Tuveson, D.A. (2019). IL1-induced JAK/STAT signaling is antagonized by TGFβ to shape CAF heterogeneity in pancreatic ductal adenocarcinoma. Cancer Discov. 9, 282–301.

38. Dominguez, C.X., Müller, S., Keerthivasan, S., Koeppen, H., Hung, J., Gierke, S., Breart, B., Foreman, O., Bainbridge, T.W., Castiglioni, A., et al. (2020). Single-cell RNA sequencing reveals stromal evolution into LRRC15+ myofibroblasts as a determinant of patient response to cancer immunotherapy. Cancer Discov. 10, 232–253.

39. Krishnamurty, A.T., Shyer, J.A., Thai, M., Gandham, V., Buechler, M.B., Yang, Y.A., Pradhan, R.N., Wang, A.W., Sanchez, P.L., Qu, Y., et al. (2022). LRRC15+ myofibroblasts dictate the stromal setpoint to suppress tumour immunity. Nature 611, 148–154.

40. Bhattacharjee, S., Hamberger, F., Ravichandra, A., Miller, M., Nair, A., Affo, S., Filliol, A., Chin, L., Savage, T.M., Yin, D., et al. (2021). Tumor restriction by type I collagen opposes tumor-promoting effects of cancer-associated fibroblasts. J. Clin. Invest. 131. 10.1172/JCI146987.

41. Maeda, K., Enomoto, A., Hara, A., Asai, N., Kobayashi, T., Horinouchi, A., Maruyama, S., Ishikawa, Y., Nishiyama, T., Kiyoi, H., et al. (2016). Identification of Meflin as a potential marker for mesenchymal stromal cells. Sci. Rep. 6, 22288.

42. Buechler, M.B., Pradhan, R.N., Krishnamurty, A.T., Cox, C., Calviello, A.K., Wang, A.W., Yang, Y.A., Tam, L., Caothien, R., Roose-Girma, M., et al. (2021). Cross-tissue organization of the fibroblast lineage. Nature 593, 575–579.

43. Lin, H., Li, N., He, H., Ying, Y., Sunkara, S., Luo, L., Lv, N., Huang, D., and Luo, Z. (2015). AMPK inhibits the stimulatory effects of TGF-β on Smad2/3 activity, cell migration, and epithelial-to-mesenchymal transition. Mol. Pharmacol. 88, 1062–1071.

44. Schug, Z.T., Peck, B., Jones, D.T., Zhang, Q., Grosskurth, S., Alam, I.S., Goodwin, L.M., Smethurst, E., Mason, S., Blyth, K., et al. (2015). Acetyl-CoA synthetase 2 promotes acetate utilization and maintains cancer cell growth under metabolic stress. Cancer Cell 27, 57–71.

45. Sahai, E., Astsaturov, I., Cukierman, E., DeNardo, D.G., Egeblad, M., Evans, R.M., Fearon, D., Greten, F.R., Hingorani, S.R., Hunter, T., et al. (2020). A framework for advancing our understanding of cancer-associated fibroblasts. Nat. Rev. Cancer 20, 174–186.

46. Chen, Y., McAndrews, K.M., and Kalluri, R. (2021). Clinical and therapeutic relevance of cancer-associated fibroblasts. Nat. Rev. Clin. Oncol. 18, 792–804.

47. Özdemir, B.C., Pentcheva-Hoang, T., Carstens, J.L., Zheng, X., Wu, C.-C., Simpson, T.R., Laklai, H., Sugimoto, H., Kahlert, C., Novitskiy, S.V., et al. (2014). Depletion of carcinoma-associated fibroblasts and fibrosis induces immunosuppression and accelerates pancreas cancer with reduced survival. Cancer Cell 25, 719–734.

48. Bangi, E., Ang, C., Smibert, P., Uzilov, A.V., Teague, A.G., Antipin, Y., Chen, R., Hecht, C., Gruszczynski, N., and Yon, W.J. (2019). A personalized platform identifies trametinib plus zoledronate for a patient with KRAS-mutant metastatic colorectal cancer. Science advances 5, eaav6528.

49. Herlemann, D.P., Labrenz, M., Jurgens, K., Bertilsson, S., Waniek, J.J., and Andersson, A.F. (2011). Transitions in bacterial communities along the 2000 km salinity gradient of the Baltic Sea. ISME J. 5, 1571–1579.

50. Shimizu, Y., Yamamura, R., Yokoi, Y., Ayabe, T., Ukawa, S., Nakamura, K., Okada, E., Imae, A., Nakagawa, T., Tamakoshi, A., et al. (2023). Shorter sleep time relates to lower human defensin 5 secretion and compositional disturbance of the intestinal microbiota accompanied by decreased short-chain fatty acid production. Gut Microbes 15, 2190306.

51. Bolyen, E., Rideout, J.R., Dillon, M.R., Bokulich, N.A., Abnet, C.C., Al-Ghalith, G.A., Alexander, H., Alm, E.J., Arumugam, M., Asnicar, F., et al. (2019). Author Correction: Reproducible, interactive, scalable and extensible microbiome data science using QIIME 2. Nat. Biotechnol. 37, 1091.

52. Lin, H., and Peddada, S.D. (2020). Analysis of compositions of microbiomes with bias correction. Nat. Commun. 11, 3514.

53. Mallick, H., Rahnavard, A., McIver, L.J., Ma, S., Zhang, Y., Nguyen, L.H., Tickle, T.L., Weingart, G., Ren, B., Schwager, E.H., et al. (2021). Multivariable association discovery in population-scale meta-omics studies. PLoS Comput. Biol. 17, e1009442.

54. Suzuki, T.A., Tanja, A.-S., Waters, J.L., Jakob, D., Vu, D.L., Ballinger, M.A., Di Rienzi, S.C., Chang, H., de Araujo, I.E., Tyakht, A.V., et al. (2025). Selection and transmission of the gut microbiome alone can shift mammalian behavior. Nat. Commun. 16, 9482.

55. Song, I., Yang, J., Saito, M., Hartanto, T., Nakayama, Y., Ichinohe, T., and Fukuda, S. (2024). Prebiotic inulin ameliorates SARS-CoV-2 infection in hamsters by modulating the gut microbiome. Npj Sci. Food 8, 18.

56. Matoba, K., Murakami, M., Fujita, E., Jin, S., Ogasawara, R., Matoba, T., Takeuchi, A., Haga, S., Ozaki, M., and Hyodoh, H. (2022). The usefulness of measuring n-butyric acid concentration as a new indicator of blood decomposition in forensic autopsy. Leg. Med. (Tokyo) 57, 102071.

57. Kato, T., Jenkins, R.P., Derzsi, S., Tozluoglu, M., Rullan, A., Hooper, S., Chaleil, R.A.G., Joyce, H., Fu, X., Thavaraj, S., et al. (2023). Interplay of adherens junctions and matrix proteolysis determines the invasive pattern and growth of squamous cell carcinoma. Elife 12. 10.7554/eLife.76520.

