## Supplemental Figures for "Co-targeting an AMPK–MAPK axis reprograms fibroblasts and suppresses PDAC"

### SUPPLEMENTAL INFORMATION

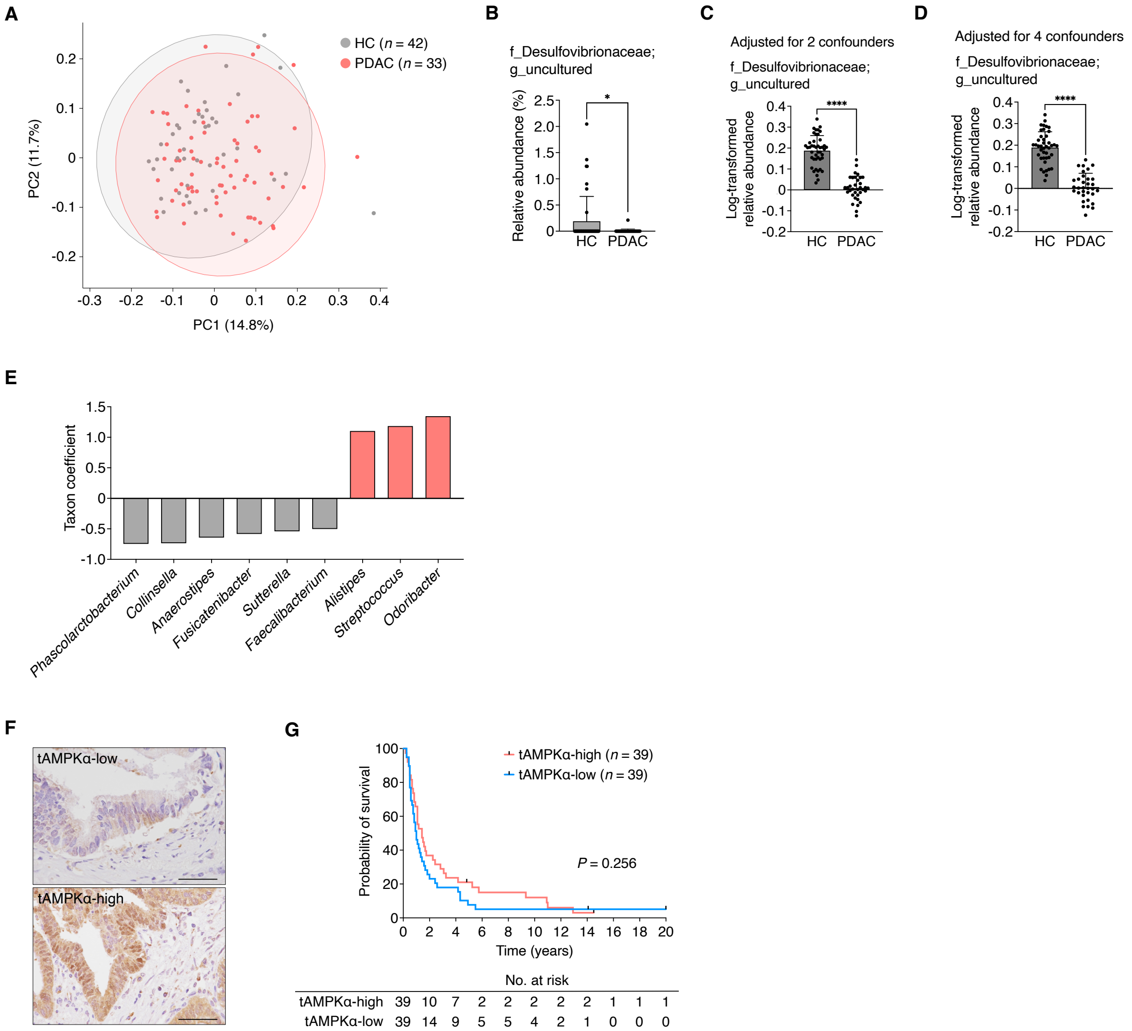


**Figure S1. Reduced abundance of AA-producing bacteria in patients with PDAC and lack of association between tAMPKα expression and prognosis, related to Figure 1**

**A,** Overall gut microbial community structure does not differ significantly between HCs ($n$ = 42) and patients with PDAC ($n$ = 33), as assessed by PCoA of weighted UniFrac distances derived from fecal 16S rRNA gene sequencing data and analyzed using PERMANOVA ($P$ = 0.080).

**B–D,** An uncultured AA-producing genus within the Desulfovibrionaceae family is significantly depleted in patients with PDAC compared with HCs.

**B,** Uncultured genus with decreased relative abundance in patients with PDAC, identified using ANCOM-BC.

**C–D,** Associations between the uncultured Desulfovibrionaceae genus and disease status assessed using MaAsLin2, adjusted for age and sex (**C**), and additionally for BMI and T2DM (**D**).

**E,** Patients with PDAC exhibit a compositional shift in the gut microbiome characterized by enrichment of PDAC-associated taxa and depletion of taxa enriched in HCs. Taxon coefficients derived from NB analysis in human samples, in which negative coefficients indicate taxa enriched in HCs (denominator taxa) and positive coefficients indicate taxa enriched in patients with PDAC (numerator taxa).

**F,** Representative images of tAMPKα IHC in PDAC tumor samples, illustrating low (tAMPKα-low) and high (tAMPKα-high) expression patterns. Scale bars, 50 μm.

**G,** Total AMPKα expression in PDAC tumors does not correlate with overall survival. Kaplan–Meier analysis of overall survival in 78 patients with PDAC who did not receive postoperative chemotherapy, stratified into tAMPKα-high ($n$ = 39) and tAMPKα-low ($n$ = 39) groups based on immunohistochemical scoring of TMA cores ($P$ = 0.256).

Each dot represents an individual human subject. Statistical significance was assessed using a two-sided Student's $t$-test (**B, G**), multi-variable linear models using MaAsLin2 (**C–D**) and the log-rank test (**G**). $^{*}P < 0.05$, $^{****}P < 0.0001$. Data are presented as the mean ± SD.

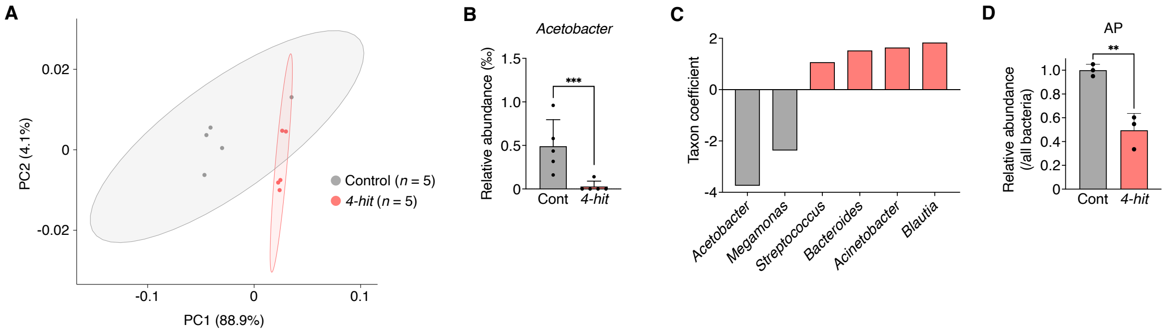


**Figure S2. Reduced abundance of AA-producing bacteria in a *Drosophila* model recapitulating core genetic alterations of PDAC, related to Figure 2**

**A,** Overall gut microbial community structure does not differ significantly between control and *4-hit* larvae, as assessed by PCoA of weighted UniFrac distances derived from midgut and hindgut samples (PERMANOVA, $P = 0.117$).

**B,** The relative abundance of *Acetobacter* is reduced in *4-hit* larvae compared with controls.

**C,** NB analysis reveals a trend toward a compositional shift in the gut microbiota of *4-hit* larvae, characterized by the enrichment of *4-hit*-associated taxa and the depletion of control-enriched taxa, including *Acetobacter*.

**D,** The abundance of the AA-producing bacterium AP is significantly reduced in *4-hit* larvae compared with controls, as quantified by qPCR and normalized to total bacterial DNA.

Each dot represents a single sample of pooled midgut and hindgut tissues from 10 fly larvae (**A–B**) or data from an independent experiment (**D**). Statistical significance was assessed using a two-sided Student's *t*-test (**B, D**). $^{**}P < 0.01$, $^{***}P < 0.001$. Data are presented as the mean ± SD.

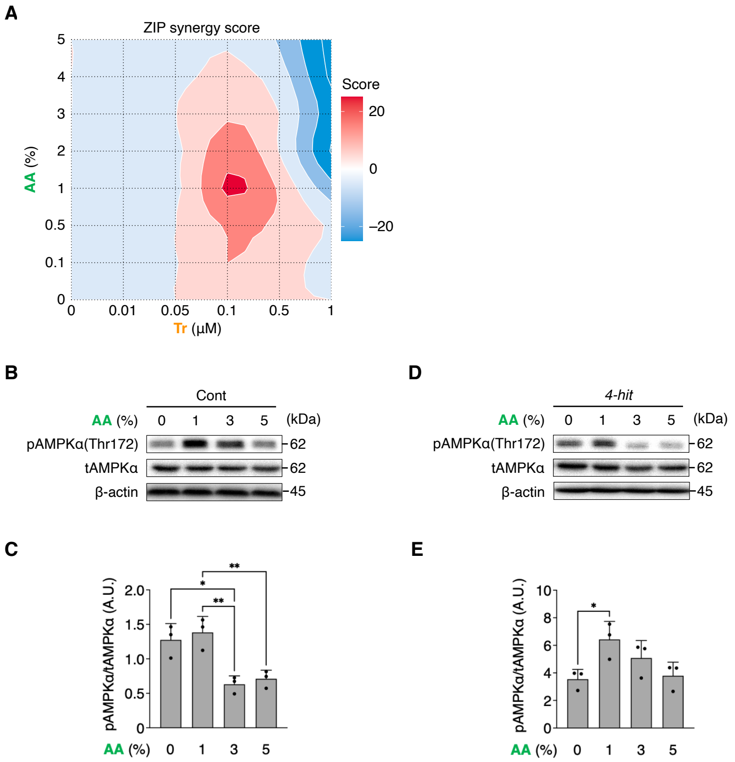


**Figure S3. AA enhances AMPK phosphorylation in *Drosophila*, related to Figure 2**

**A,** ZIP synergy analysis identifies the combination of 0.1 μM Tr and 1% AA as yielding the greatest survival benefit in *4-hit* flies.

**B–E,** AA exposure induces AMPKα phosphorylation in both control *Ser-gal4* larvae (**B**) and *4-hit* larvae (**D**), although this activation is attenuated in *4-hit* larvae. β-actin served as a loading control. Densitometric quantification of the pAMPKα/tAMPKα ratio is presented for control (**C**) and *4-hit* (**E**) larvae.

Each dot represents an independent experiment (**C, E**). Statistical significance was assessed using one-way ANOVA followed by Tukey's multiple comparisons test (**D, F**). $^{*}P < 0.05$, $^{**}P < 0.01$, $^{***}P < 0.001$, $^{****}P < 0.0001$. Data are presented as the mean ± SD. A.U., arbitrary units.

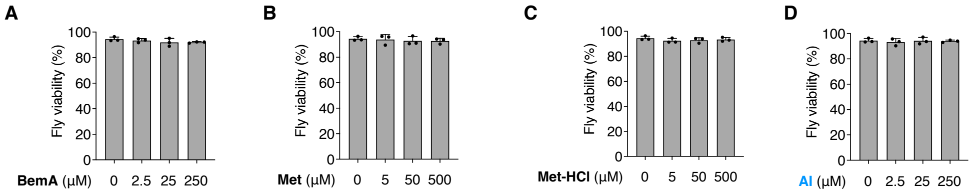


**Figure S4 Determination of the MTDs of AMPK activators in *Drosophila*, related to Figure 3**

**A–D,** All four tested AMPK activators were well tolerated by non-transgenic control (*w*–) flies, thereby establishing their MTDs. Viability is shown for control flies reared on medium containing increasing concentrations of BemA (0–250 μM) (**A**), Met (0–500 μM) (**B**), Met-HCl (0–500 μM) (**C**), and AI (0–250 μM) (**D**). These assays confirm that the concentrations used in subsequent experiments do not impair the survival of wild-type flies.

Each dot represents an independent experiment. Data are presented as the mean ± SD.

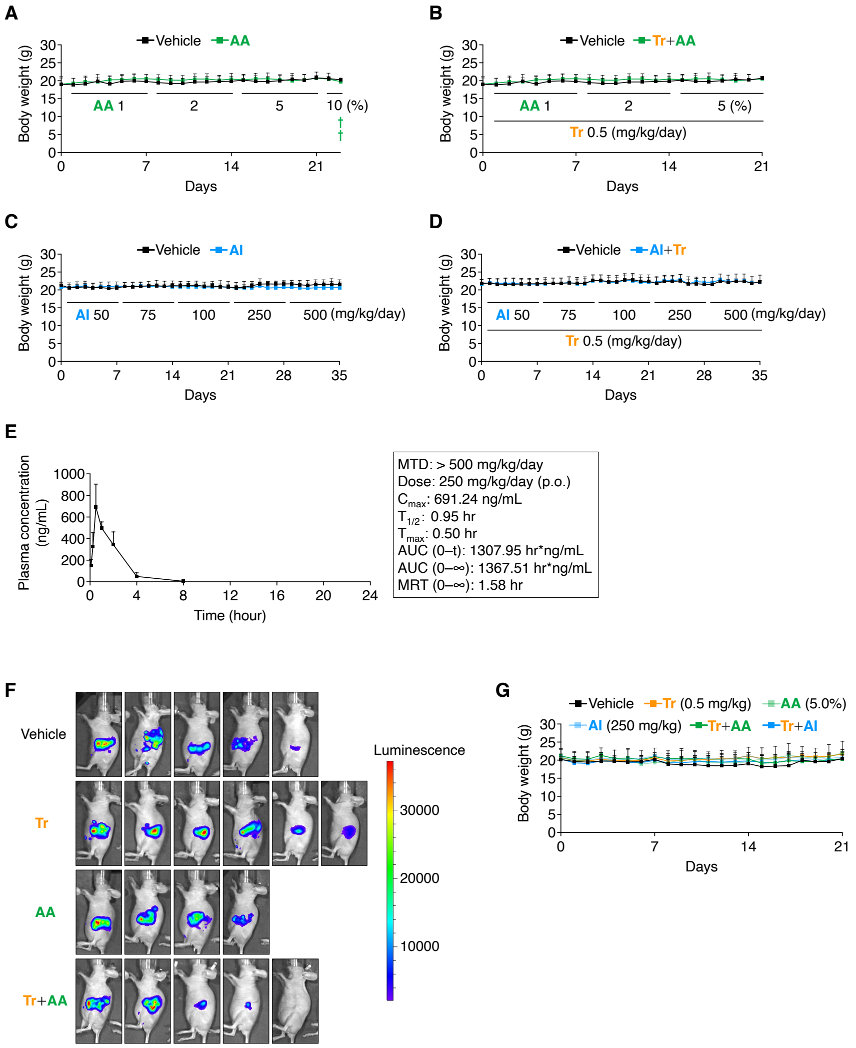


**Figure S5 Determination of the optimal dosing, pharmacokinetic, and safety profiles of compounds in mice, related to Figure 4**

**A–B,** Monitoring of body weight established 5% AA as the MTD in female BALB/c-*nu/nu* mice. Shown are body weight changes in mice treated with AA (1%–10%) alone (**A**) or in combination with Tr (0.5 mg/kg) (**B**) over a 21-day period. †, denotes the death of one mouse.

**C–D,** Monitoring of body weight demonstrates that AI is well tolerated at doses up to 500 mg/kg, both as monotherapy (**C**) and in combination with Tr (**D**), in female BALB/c-*nu*/*nu* mice.

**E,** PK analysis of AI following a single oral dose (250 mg/kg) in female BALB/c-*nu*/*nu* mice. The plasma concentration–time profile and key PK parameters, including maximum plasma concentration ($C_{max}$), half-life ($T_{1/2}$), time to maximum plasma concentration ($T_{max}$), area under the curve and mean residence time, are presented.

**F,** Individual tumor growth, monitored by bioluminescence imaging, is shown for each treatment cohort.

**G,** Tr+AI does not induce significant body weight loss compared with vehicle, indicating good *in vivo* tolerability. Body weight changes are presented for mice treated with vehicle, Tr (0.5 mg/kg), AA (5%), AI (250 mg/kg) or their combinations. †, denotes the death of one mouse.

Data are presented as the mean ± SD.

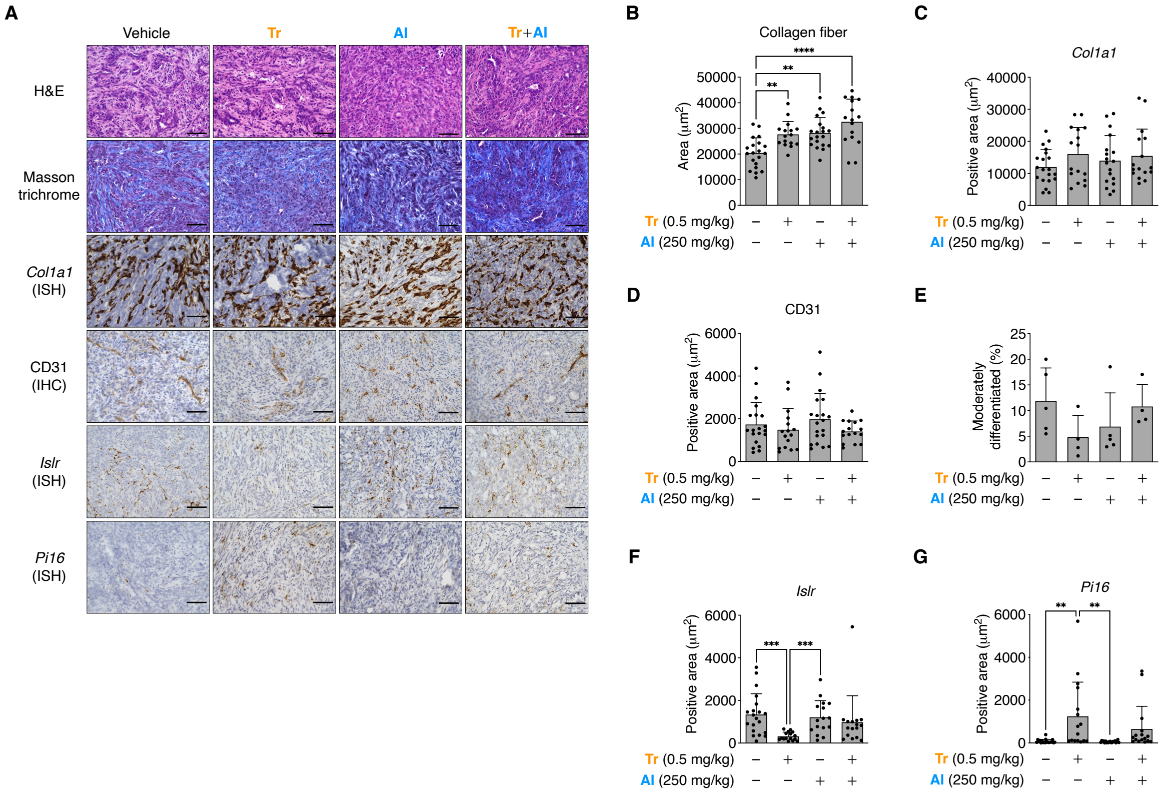


**Figure S6 Co-targeting MEK and AMPK selectively reshapes the stromal microenvironment, related to Figure 5**

**A,** Representative histological images of allograft tumors harvested on day 10. Tumor sections were stained with hematoxylin and eosin, Masson trichrome, *Col1a1* by ISH, CD31 by IHC, *Islr* by ISH and *Pi16* by ISH. Scale bars, 100 µm.

**B–G,** Co-targeting MEK and AMPK selectively enhances collagen deposition while producing minimal effects on tumor vasculature or tumor differentiation, and modulates specific stromal marker expression. Quantification of the Masson's trichrome-positive collagen area (**B**), *Col1a1* expression area (**C**), CD31-positive endothelial area (**D**), *Pi16* expression area (**E**), *Islr* expression area (**F**) and the proportion of moderately differentiated tumor regions (**G**).

Each dot represents a single microscopic field (**B–G**), with four randomly selected regions imaged and evaluated per allograft. Statistical significance was assessed using one-way ANOVA followed by Tukey's multiple comparisons test (**B–E, G**), and by the Kruskal–Wallis test followed by Dunn's multiple comparisons test (**F**). $^{**}P < 0.01$, $^{***}P < 0.001$, $^{****}P < 0.0001$. Data are presented as the mean ± SD.

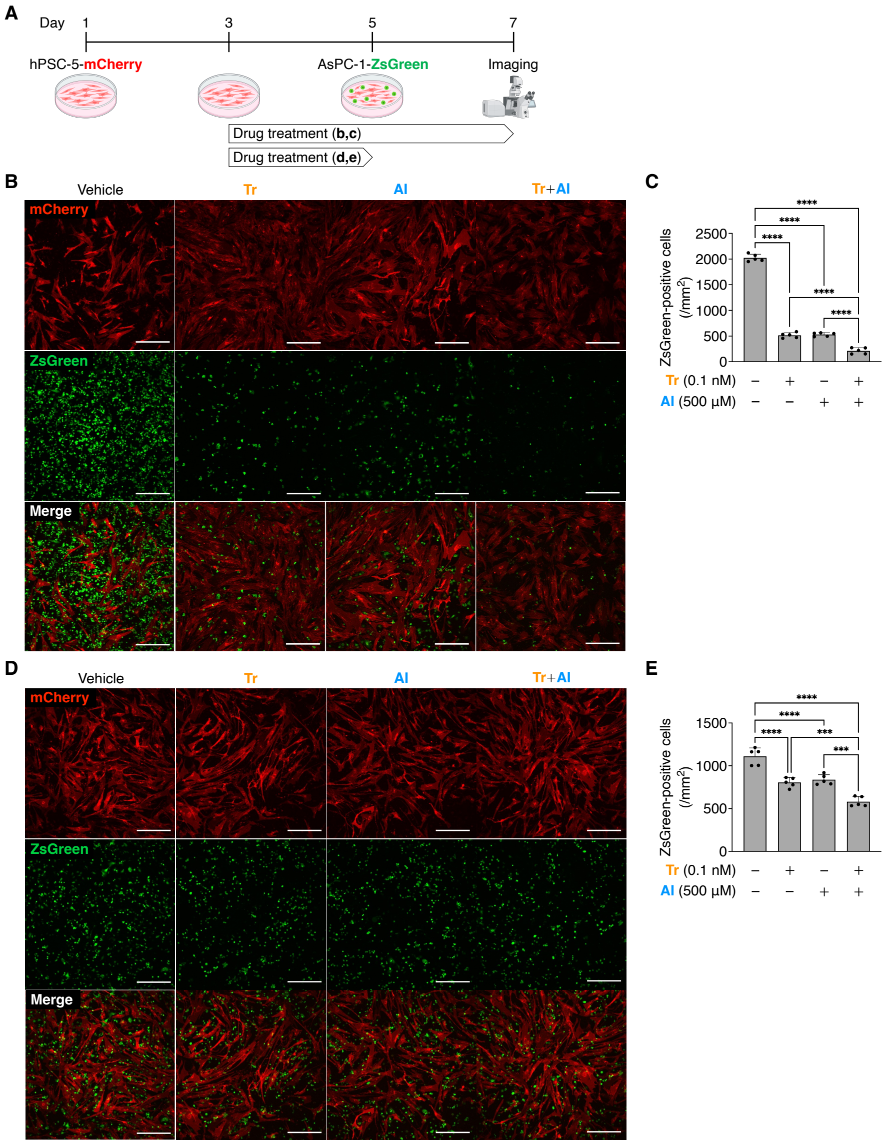


**Figure S7 Dual modulation of MAPK and AMPK suppresses CAF-driven PDAC cell proliferation, related to Figure 6**

**A,** Schematic of the co-culture experimental design. hPSC-5-mCherry CAFs were seeded on collagen I-coated glass-bottom dishes and cultured for 48 h, followed by treatment with DMSO, Tr (0.1 nM), AI (500 μM) or the Tr+AI combination for 48 h. AsPC-1-ZsGreen cancer cells were then added to the same wells and co-cultured for an additional 48 h, either with or without continued drug exposure. Fluorescence images were acquired using a confocal microscope under identical acquisition settings, and AsPC-1-ZsGreen cell numbers were quantified.

**B–C,** Continuous drug exposure significantly suppresses AsPC-1 proliferation, with the Tr + AI combination exhibiting the strongest inhibitory effect. Representative fluorescence images (**B**) and quantification of AsPC-1-ZsGreen cells (green) (**C**) in co-culture with hPSC-5-mCherry CAFs (red) (B). CAFs were pre-treated with DMSO, AI (500 μM), Tr (0.1 nM) or Tr+AI for 48 h, after which AsPC-1 cells were added and co-cultured for an additional 48 h under continued drug exposure. Scale bars, 400 μm.

**D–E,** Drug withdrawal at the time of AsPC-1 seeding attenuates, but does not completely abrogate, the suppressive effect on cancer cell proliferation. Representative fluorescence images (**D**) and quantification of AsPC-1 cell numbers (**E**) in co-cultures in which CAFs were pre-treated as in (**B–C**), followed by drug removal at the time of AsPC-1 seeding and subsequent 48-h co-culture in drug-free medium. Scale bars, 400 μm.

Each dot represents an independent experiment (**C, E**). Statistical significance was assessed using one-way ANOVA followed by Tukey’s multiple comparisons test (**C, E**). $^{***}P < 0.001$, $^{****}P < 0.0001$. Data are presented as the mean ± SD.
