## Supplemental Tables for "Co-targeting an AMPK–MAPK axis reprograms fibroblasts and suppresses PDAC"

**Table S1**
**Characteristics of the study participants, related to Figure 1**

| | HC ($n$ = 42) | | PDAC ($n$ = 33) | | *P*-value |
|---|---|---|---|---|---|
| Age, years | 66.4 | (9.3) | 68.2 | (8.8) | 0.417 |
| Gender | | | | | |
| Male | 23 | (54.8) | 20 | (60.6) | 0.785 |
| Female | 19 | (45.2) | 13 | (39.4) | |
| BMI, kg/m$^2$ | 22.8 | (3.1) | 21.5 | (3.2) | 0.062 |
| Diabetes melitus | 5 | (11.9) | 7 | (21.2) | 0.439 |
| Tumor location | | | | | |
| Head | – | | 15 | (45.5) | – |
| Body/Tail | – | | 18 | (54.5) | – |
| Stage[#] | | | | | |
| I/II | – | | 26 | (78.8) | – |
| III/IV | – | | 7 | (21.2) | – |
| cT (tumor size) | | | | | |
| 1+2 | – | | 21 | (63.6) | – |
| 3+4 | – | | 12 | (36.4) | – |
| cN (regional lymph node metastasis) | | | | | |
| Negative | – | | 28 | (84.8) | – |
| Positive | – | | 5 | (15.2) | – |
| cM (distant metastasis) | | | | | |
| Negative | – | | 24 | (72.7) | – |
| Positive | – | | 9 | (27.3) | – |

Abbreviations: BMI, body mass index; HC, healthy control; PDAC, pancreatic ductal adenocarcinoma; SD, standard deviation. Values are presented as mean ± standard deviation, or the number (%) of participants in that category. [#]The staging of tumors was determined based on the American Joint Committee on Cancer/Union for International Cancer Control (AJCC/UICC) 8th edition guidelines. *P*-values were calculated using one-way analysis of variance (ANOVA).

**Table S2**

**Fecal levels of nine organic acids in HC and PDAC, related to Figure 1**

| | HC (*n* = 42) | | PDAC (*n* = 33) | | *P*-value |
|---|---|---|---|---|---|
| AA (µmol/g feces) | | | | | |
| Crude mean (SD) | 271.8 | (133.8) | 165.8 | (104.1) | < 0.001 |
| Adjusted LSM (SE), model 1 | 268.8 | (18.2) | 157.1 | (20.8) | < 0.001 |
| Adjusted LSM (SE), model 2 | 237.9 | (23.1) | 133.0 | (23.1) | < 0.001 |
| BA (µmol/g feces) | | | | | |
| Crude mean (SD) | 47.9 | (42.4) | 23.7 | (22.1) | 0.004 |
| Adjusted LSM (SE), model 1 | 47.2 | (5.3) | 21.5 | (6.1) | 0.002 |
| Adjusted LSM (SE), model 2 | 52.8 | (6.9) | 25.2 | (6.9) | 0.002 |
| FA (µmol/g feces) | | | | | |
| Crude mean (SD) | 2.0 | (0.5) | 1.8 | (0.6) | 0.126 |
| Adjusted LSM (SE), model 1 | 2.0 | (0.1) | 1.8 | (0.1) | 0.150 |
| Adjusted LSM (SE), model 2 | 2.0 | (0.1) | 1.7 | (0.1) | 0.059 |
| IBA (µmol/g feces) | | | | | |
| Crude mean (SD) | 6.3 | (3.8) | 6.5 | (4.1) | 0.773 |
| Adjusted LSM (SE), model 1 | 6.3 | (0.6) | 6.3 | (0.7) | 0.990 |
| Adjusted LSM (SE), model 2 | 6.9 | (0.8) | 6.3 | (0.8) | 0.563 |
| IVA (µmol/g feces) | | | | | |
| Crude mean (SD) | 7.5 | (4.8) | 7.5 | (4.5) | 0.996 |
| Adjusted LSM (SE), model 1 | 7.5 | (0.7) | 7.2 | (0.8) | 0.779 |
| Adjusted LSM (SE), model 2 | 9.0 | (0.9) | 7.9 | (0.9) | 0.308 |
| LA (µmol/g feces) | | | | | |
| Crude mean (SD) | 3.0 | (9.8) | 31.6 | (167.9) | 0.274 |
| Adjusted LSM (SE), model 1 | 4.1 | (17.2) | 35.1 | (19.8) | 0.241 |
| Adjusted LSM (SE), model 2 | 28.6 | (21.6) | 59.3 | (21.7) | 0.250 |
| PA (µmol/g feces) | | | | | |
| Crude mean (SD) | 94.8 | (61.7) | 49.3 | (37.5) | < 0.001 |
| Adjusted LSM (SE), model 1 | 93.1 | (7.8) | 45.6 | (8.8) | < 0.001 |
| Adjusted LSM (SE), model 2 | 84.2 | (10.0) | 37.8 | (10.0) | < 0.001 |
| SA (µmol/g feces) | | | | | |
| Crude mean (SD) | 2.7 | (7.6) | 1.3 | (1.7) | 0.307 |
| Adjusted LSM (SE), model 1 | 2.6 | (0.9) | 1.1 | (1.0) | 0.246 |
| Adjusted LSM (SE), model 2 | 2.3 | (1.2) | 0.6 | (1.2) | 0.231 |

**Table S2 (continued)**

| | | | | | |
|---|---|---|---|---|---|
| VA (µmol/g feces) | | | | | |
| Crude mean (SD) | 8.0 | (7.5) | 5.3 | (5.6) | 0.096 |
| Adjusted LSM (SE), model 1 | 7.9 | (1.0) | 5.0 | (1.2) | 0.070 |
| Adjusted LSM (SE), model 2 | 8.3 | (1.4) | 5.3 | (1.4) | 0.072 |

Abbreviations: AA, acetic acid; BA, butyric acid; FA, formic acid; HC, healthy control; IBA, isobutyric acid; IVA, isovaleric acid; LA, lactic acid; LSM, least square mean; PDAC, pancreatic ducatal adenocarcinoma; PA, propionic acid; SA, succinic acid; SCFAs, short-chain fatty acids; SD, standard deviation; SE, standard error; VA, valeric acid. *P*-values were calculated using a two-sided Student's *t*-test. In addition, analysis of covariance (ANCOVA) was performed to adjust for potential confounding factors. Model 1 was adjusted for age and sex, while Model 2 included the covariates from Model 1 along with body mass index (BMI) and diabetes mellitus (DM) status.

**Table S3**

**Clinical characteristics of TMA cases stratified by phospho-AMPK expression status, related to Figure 1**

| | pAMPKα negative (*n* = 41) | | pAMPKα positive (*n* = 37) | | *P*-value |
|---|---|---|---|---|---|
| Age, years | 67.6 | (8.7) | 65.9 | (9.5) | 0.413 |
| Gender | | | | | |
| Male | 26 | (63.4) | 24 | (64.9) | 1.000 |
| Female | 15 | (36.6) | 13 | (35.1) | |
| Tumor location | | | | | |
| Head | 25 | (61.0) | 26 | (70.3) | 0.533 |
| Body/Tail | 16 | (39.0) | 11 | (29.7) | |
| Pathological stage# | | | | | |
| I/II | 8 | (19.5) | 12 | (32.4) | 0.296 |
| III/IV | 33 | (80.5) | 25 | (67.6) | |
| pT (tumor size) | | | | | |
| 1+2 | 27 | (65.9) | 28 | (75.7) | 0.483 |
| 3+4 | 14 | (34.1) | 9 | (24.3) | |
| pN (regional lymph node metastasis) | | | | | |
| Negative | 9 | (22.0) | 12 | (32.4) | 0.432 |
| Positive | 32 | (78.0) | 25 | (67.6) | |
| pM (distant metastasis) | | | | | |
| Negative | 36 | (87.8) | 35 | (94.6) | 0.515 |
| Positive | 5 | (12.2) | 2 | (5.4) | |
| Differentiation | | | | | |
| Well to moderately | 34 | (82.9) | 35 | (94.6) | 0.209 |
| Poorly and other | 7 | (17.1) | 2 | (5.4) | |

Abbreviations: pAMPKα, phospho-AMP-activated protein kinase α; PDAC, pancreatic ductal adenocarcinoma; SD, standard deviation. Values are presented as mean (SD), or the number (%) of participants in that category. #The staging of tumors was determined based on the American Joint Committee on Cancer/Union for International Cancer Control (AJCC/UICC) 8th edition guidelines. *P*-values were calculated using one-way analysis of variance (ANOVA).

**Table S4**

**Prognostic factors for overall survival of PDAC patients, related to Figure 1**

| | OS (years) | | |
|---|---|---|---|
| | HRs of death | (95% CI) | *P*-value |
| Phosho-AMPKα (positive vs. negative), model 1 | 1.66 | (1.03 – 2.65) | 0.037 |
| Phosho-AMPKα (positive vs. negative), model 2 | 1.62 | (0.99 – 2.65) | 0.057 |

Abbreviations: CI, confidence interval; HRs, hazard ratios; OS, overall survival; PDAC, pancreatic ductal adenocarcinoma. HRs, 95% CIs, and *P*-values were calculated using the Cox proportional hazards model. Model 1 was not adjusted for any confounding factors; model 2 was adjusted for patients' age and sex.

**Table S5**

**Clinical characteristics of TMA cases stratified by total AMPK expression status, related to Figure 1**

| | tAMPKα negative (*n* = 39) | | tAMPKα positive (*n* = 39) | | *P*-value |
|---|---|---|---|---|---|
| Age, years | 66.1 | (9.6) | 67.4 | (8.6) | 0.528 |
| Gender | | | | | |
| Male | 28 | (71.8) | 22 | (56.4) | 0.238 |
| Female | 11 | (28.2) | 17 | (43.6) | |
| Tumor location | | | | | |
| Head | 24 | (61.5) | 27 | (69.2) | 0.634 |
| Body/Tail | 15 | (38.5) | 12 | (30.8) | |
| Pathological stage# | | | | | |
| I/II | 8 | (20.5) | 12 | (30.8) | 0.437 |
| III/IV | 31 | (79.5) | 27 | (69.2) | |
| pT (tumor size) | | | | | |
| 1+2 | 26 | (66.7) | 29 | (74.4) | 0.620 |
| 3+4 | 13 | (33.3) | 10 | (25.6) | |
| pN (regional lymph node metastasis) | | | | | |
| Negative | 9 | (23.1) | 12 | (30.8) | 0.610 |
| Positive | 30 | (76.9) | 27 | (69.2) | |
| pM (distant metastasis) | | | | | |
| Negative | 33 | (84.6) | 38 | (97.4) | 0.113 |
| Positive | 6 | (15.4) | 1 | (2.6) | |
| Differentiation | | | | | |
| Well to moderately | 34 | (87.2) | 35 | (89.7) | 1.000 |
| Poorly and other | 5 | (12.8) | 4 | (10.3) | |

Abbreviations: PDAC, pancreatic ductal adenocarcinoma; SD, standard deviation; tAMPKα, total AMP-activated protein kinase α. Values are presented as mean (SD), or the number (%) of participants in that category. #The staging of tumors was determined based on the American Joint Committee on Cancer/Union for International Cancer Control (AJCC/UICC) 8th edition guidelines. *P*-values were calculated using one-way analysis of variance (ANOVA).

**Table S6**

**Prognostic factors for overall survival of PDAC patients, related to Figure 1**

| | OS (years) | | |
|---|---|---|---|
| | HRs of death | (95% CI) | *P*-value |
| Total AMPKα (positive vs. negative), model 1 | 0.79 | (0.50 – 1.25) | 0.308 |
| Total AMPKα (positive vs. negative), model 2 | 0.74 | (0.45 – 1.23) | 0.246 |

Abbreviations: CI, confidence interval; HRs, hazard ratios; OS, overall survival; PDAC, pancreatic ductal adenocarcinoma. HRs, 95% CIs, and *P*-values were calculated using the Cox proportional hazards model. Model 1 was not adjusted for any confounding factors; model 2 was adjusted for patients' age and sex.
